# Immune cells regulate circulating adipocyte extracellular vesicle levels in response to metabolic shifts

**DOI:** 10.1101/2025.07.11.664236

**Authors:** Snigdha Tiash, Marjori Russo, Yun-Ling Pai, Wentong Jia, Rachael L. Field, Terri Pietka, Nada A. Abumrad, Gordon I. Smith, Dmitri Samovski, Samuel Klein, Jonathan R. Brestoff, Clair Crewe

## Abstract

Extracellular vesicles (EVs) are now recognized as potent mediators of intercellular and inter-organ signaling and implicated in the pathogenesis of obesity and its associated comorbidities such as diabetes, cancer, cardiovascular disease, and neurodegeneration. Despite a surge of new functional information about EVs, we still lack a basic understanding of how endogenous EV levels are controlled to regulate inter-organ signaling. New flow cytometry technology has allowed us to study the regulation of circulating, endogenous EVs from metabolically relevant cell types like adipocytes. From this, we provide evidence for a paradigm of EV regulation where tissue resident immune cells, predominantly macrophages, clear EVs released by local tissue cells or EVs entering the tissue from circulation, an activity that determines circulating EV levels. In obesity, EV uptake by adipose tissue immune cells is reduced, concomitant with increased circulating adipocyte-specific EVs (adipoEVs) and reduced EV clearance rates. AdipoEVs are significantly elevated in mouse circulation from one day to 20 weeks of high-fat feeding. In humans we found that adipocyte EV levels negatively correlate with whole-body and liver insulin sensitivity and are not associated with adipose mass. This work suggests that tissue resident immune cells act as a gatekeeper for tissue EV entry into circulation and are thereby a major regulator of inter-organ EV signaling.

## INTRODUCTION

Extracellular vesicles (EVs) are powerful modes of inter-cellular communication that have recently been shown to participate in the progression of cardiometabolic diseases^1,2^. The ability of these nano-sized vesicles to signal to a recipient cell is owed to the diverse cargo they carry from various RNA biotypes, to signaling proteins, metabolites and lipids^3^. EVs can signal to other cells in the tissue or may enter circulation and modulate the function of cells in other organs. Adipocyte-derived EVs (adipoEVs) have emerged as robust regulators of metabolism in both health and disease. AdipoEVs are replete in microRNAs and can regulate function of the liver, endothelium, pancreas, and brain^4–10^. AdipoEVs are also laden with lipids that can fuel cancer cells or exercising muscle^11–13^. In the context of obesity, adipoEVs are generally considered pro-pathogenic, by transmitting adverse phenotypes like insulin resistance, inflammation, and fibrosis from the adipose tissue to other organs, particularly the liver^14–16^. In contrast, adipocyte EVs contain active adiponectin which has insulin-sensitizing effects^17^. Furthermore, adipocytes can package pieces of damaged mitochondria into EVs (EV-mito), which are cardioprotective^18^.

Despite the growing body of information about EVs in metabolic regulation, there are still fundamental aspects of EV physiology that are unknown. The most critical is our lack of knowledge about how endogenously produced EV levels are regulated *in vivo.* To date, we have no information about the concentration of metabolic cell-type specific EVs in circulation and what factors determine their levels. Similarly, whether cell-type specific EV levels are differentially regulated under various physiological conditions remains unknown. As circulation is the conduit by which EVs signal between organs, the ability to quantify cell-type specific EVs in blood can provide important information about the physiological context in which these EVs signal. The dominant reason for this gap in knowledge is, until recently, there has been no instrumentation able to detect fluorescent markers on single EVs in the size range that has been used for most *in vivo* functional studies (<200nm), outside of low throughput imaging techniques. Others have used computational approaches to estimate the relative abundance of plasma EVs from specific cells based on RNA cargo^19^. Although this approach is insightful, it does not provide absolute abundances, is low throughput, requires a large amount of starting plasma, and requires expertise in bioinformatics. Others have pioneered the methodology for the use of flow cytometers to detect makers for blood cell-derived large EVs (lEVs) (endothelial cells, leukocytes, and platelets; >200 nm, or the pellet of a 10,000 xg spin) in humans. Changes in these blood cell lEVs were associated with vascular insulin resistance, cardiorespiratory fitness, post prandial glucose load, hypertension, and exercise in humans^20–25^.

In this study, we took advantage of new flow cytometry technology with enhanced small particle detection that allows for multiplexed detection of markers on individual EVs down to ∼80nm in diameter. We developed a pipeline that stains EVs in just 2 µl of mouse or human plasma and purifies EVs with size exclusion chromatography (SEC) up to 96 samples at a time. This allowed us to provide the first estimate of the physiological concentration of circulating EVs from metabolically important cells, such as adipocytes, hepatocytes, and myocytes in lean and obese mice. Because adipocyte-derived EVs and EV-mitos are relevant to multiple obesity associated co-morbidities like cardiovascular diseases, cancers, and cognitive decline, we focused on the how circulating levels of these EVs are regulated. We demonstrate that total EVs and adipocyte-specific EVs (adipoEVs) are robustly regulated in response to metabolic shifts with high-fat feeding and fasting/re-feeding transitions. Macrophage-mediated clearance is responsible for most of this regulation in lean mice. In obese mice, circulating EVs, are stabilized due to slowed clearance, resulting in a greater accessibility for EVs to signal between organs. In humans, adipocyte EVs are associated with whole-body and hepatic insulin resistance. This is consistent with data in pre-clinical models that demonstrate adipoEVs can directly promote insulin resistance^14,16,26,27^. This work proposes a model of EV regulation where in healthy tissue immune cells intercept EVs before they can enter the blood resulting in low levels of interorgan signaling. In obesity, reduced EV uptake by immune cells allows tissue-specific EVs to accumulate in circulation and accelerate interorgan signaling.

## RESULTS

### Development of assays for the flow cytometric quantification of cell type-specific EVs in plasma

We used the Cytek Northern Lights spectral flow cytometer with enhanced small particle detection to quantify EVs stained with either a pan EV stain (carboxyfluorescein succinimidyl ester, CFSE) or fluorescent antibodies against cell-type specific markers. Our settings enabled detection of 100 nm polystyrene microspheres (**Fig. 1A**). Calibration of the SSC detector with Mie theory revealed the presence of particles in mouse plasma from ∼80 nm to 2 µm in diameter (**Fig. 1B**). The analysis was biased toward larger particles; however the majority of EVs are likely detected. Flow cytometers with a ∼110 nm detection limit allows for detection of 71% of EVs^28^. The total number of particles at all sizes increased in the plasma of obese mice, but also coincided with a new, larger population of particles (**Fig. 1B**). CFSE dye was used to mark all EVs. CFSE is a membrane-permeable dye that covalently conjugates a fluorophore to proteins and is previously validated as a robust EV dye for flow cytometry with minimal artifacts^29^. This reaction requires cellular esterases that are enriched in EVs, but not lipoproteins. CFSE positive EVs were increased in obese plasma ∼4 fold over that from lean mice (**Fig. 1C and D**). We can use a cocktail of EV makers and lipoprotein makers to demonstrate the clear separation of these populations (**Fig. 1E**).

**Figure 1.**
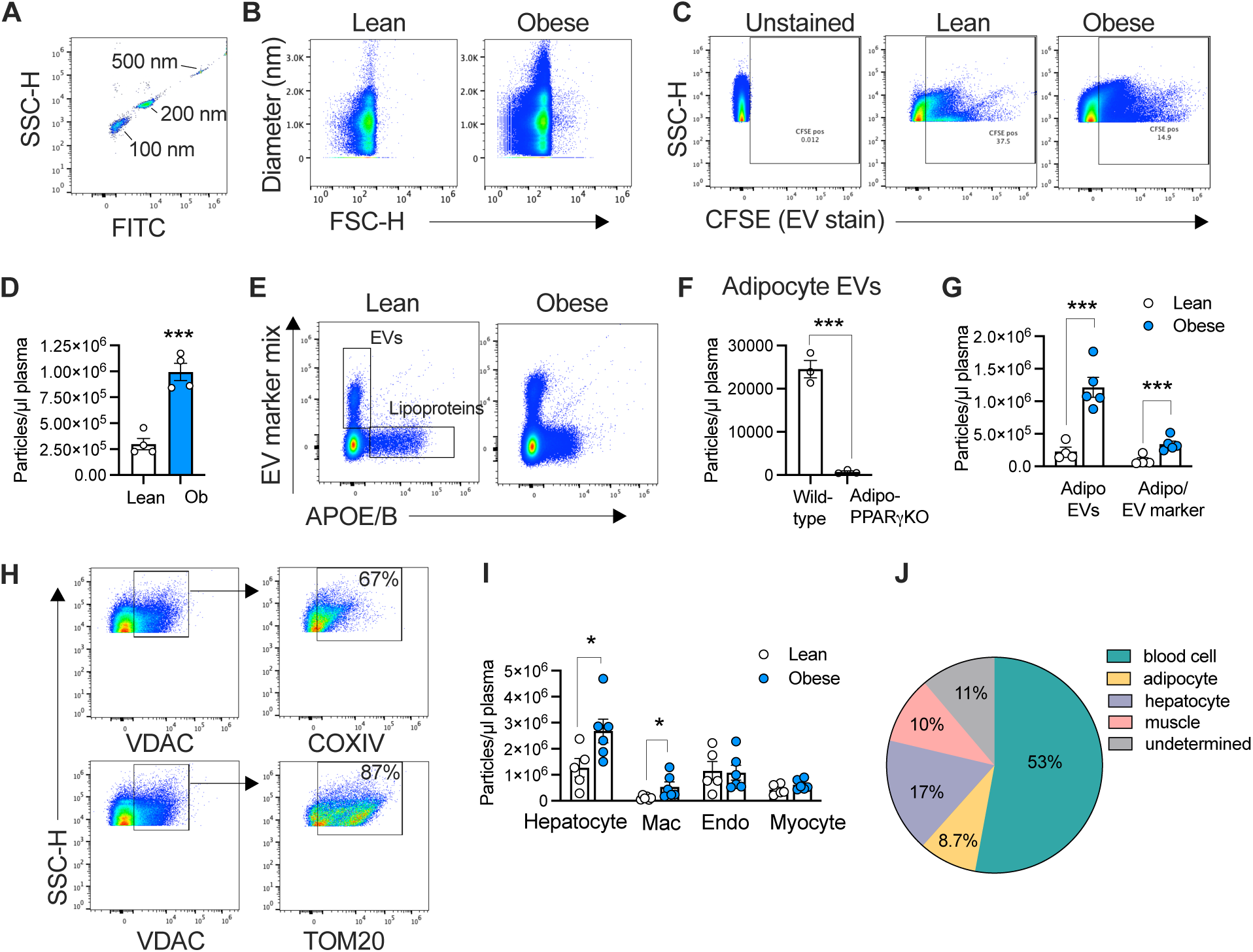
Validation of nano particle flow cytometry methodology. **A.** polystyrene microspheres of the indicated size. **B**. EVs were enriched from lean (normal chow) or obese (12 weeks high-fat diet) mouse plasma (2µl), with size exclusion chromatography (SEC) and analyzed by flow cytometry. The SSC detector was calibrated to display EV diameter **C.** CFSE stained EVs from lean and obese mice processed as in B. **D.** quantification of C **E.** A representative experiment where plasma is labeled with a cocktail of antibodies for EVs (CD63, CD81, ITGB1), or lipoproteins (APOE, APOB). EVs were enriched with SEC before flow analysis. **F.** Mouse plasma stained with an adipocyte marker mix (perilipin 1 and adiponectin). Plasma was harvested from either wild type mice or adipocyte-specific PPARγ knockout mice (Adipo-PPARγKO). **G**. Plasma stained from lean or obese (12 weeks high-fat diet). Total EVs that stain with the adipocyte marker mix (AdipoEVs) and EVs that co-stain with the both the adipocyte and EV marker mixes (Adipo/EV maker). **H.** Mitochondria isolated from mouse subcutaneous adipose tissue and stained with mitochondrial proteins: VDAC and COXIV or VDAC and TOM20. **I.** Plasma from mice fed a chow or high-fat diet (6 weeks) stained with markers for the indicated cell types. **J.** The percentage of EVs from each cell type in mouse plasma. Data are presented as mean ± s.e.m. \**P* < 0.05, *** *P* < 0.001

To develop an assay for EVs specifically from adipocytes we tested several antibodies for adipocyte-specific or enriched markers including fatty acid binding protein 4 (FABP4), hormone sensitive lipase (HSL), adiponectin (APN), and perilipin 1 (PLN1). The most specific antibodies were APN and PLN1, and when combined provided the brightest detection of adipocyte EVs in plasma. Adiponectin has previously been identified as adipocyte EV cargo^17^. We validated the specificity of our antibodies by staining plasma EVs of WT mice or adipocyte-specific PPARγ knockout mice (Adipo-PPARγKO), which do not develop adipose tissue. Adipocyte EVs (adipoEVs) were not detectable in the plasma of Adipo-PPARγKO but were present in WT plasma (**Fig. 1F**). Obese mice displayed significantly higher plasma adipocyte EVs (**Fig. 1G**). This was determined by both the total EVs with adipocyte markers and EVs that were co-stained with adipocyte makers and EV the marker mix (CD9/CD63/ITGB1; **Fig. 1G**). Because there is no universal marker that will stain all EVs, only a minority of APN/PLN1^+^ EVs also stained with the EV marker mix (Fig. 1G). EVs that contained both the adipocyte markers, and a mitochondrial marker were quantified as adipocyte-derived EV-mitos. “EV-mitos” are defined as mitochondria enclosed in an EV membrane^30^. Of the mitochondrial markers tested (COXIV, HSP60, VDAC, TOM20), VDAC provided the most robust labeling of plasma EVs at the lowest concentration of antibody. We validated the specificity of this antibody by co-staining mitochondria isolated from the subcutaneous white adipose tissue (sWAT) with VDAC and either TOM20 or COXIV. We gated on mitochondria above 500 nm in size to exclude cellular debris. 67% of VDAC^+^ mitochondria also stained for COXIV and 87% were also positive for TOM20 (**Fig. 1H**). Asialoglycoprotein receptor 1 (ASGR1) was used as a hepatocyte-specific marker as previously described^31^. EVs positive for ASGR1 and negative for CD45 were defined hepatocyte-derived EVs. Myosin Heavy Chain (MHC) was used as a marker for skeletal muscle EVs. Isotype controls confirmed that all antibodies used in this study were producing a specific signal (**Fig. S1**). At 6 weeks of high-fat feeding, male mice displayed a significant increase in, macrophage (F480^+^), and hepatocyte EVs (**Fig. 1I**). No change was detected in myocyte or endothelial cell (CD31^+^) EVs (**Fig. 1I**). In lean, healthy mice, adipocyte EVs make up 8.7% of total circulating EVs, hepatocyte EVs make up 17% and myocyte EVs make up 10% (**Fig. 1J**). The majority (53%) come from blood cells in which we included hematopoietic cells (immune cells, platelets, and red blood cells), and endothelial cells which are in direct contact with the blood. These percentages were determined by staining plasma with the brightest available fluorophore for each marker to prevent inaccurate estimates caused by differences in fluorophore brightness. Table S1 provides the physiological range of plasma EVs from adipocytes, myocytes, and hepatocytes.

### Adipocyte EVs and EV-mitos are persistently increased in circulation during obesity development

To understand the timing of EV appearance in the blood with high-fat feeding, we placed a cohort of male and female mice on either a high-fat diet or matched chow diet. Plasma samples were taken to quantify cell-specific EVs in plasma at baseline (0 days on diet), after acute high fat feeding (1-7 days of feeding), or chronic high-fat feeding (2-18 weeks of feeding). Plasma was stained with two panels of antibodies, a blood cell panel (CD45, TER119, CD31, and CD41) or an adipose tissue panel (APN/PLN1, F4/80, VDAC; **Fig. S2**). Mice gained weight on a high-fat diet as expected (**Fig. 2A**) and more lipoproteins were detected in the plasma (**Fig. 2B**). Our first cohort displayed a ∼50% reduction in total circulating EVs at the 24-hour timepoint compared to baseline regardless of their diet (chow diet, **Fig. 2C**). Zierden et. al. reported that stress-associated cortisol signaling reduces circulating EVs^32^. We found that if mice have never been handled, circulating EVs are reduced by 50% within 30 minutes of the first bleed (**Fig. 2D**), further suggesting an effect of stress on blood EV concentration. For a more accurate understanding of the acute effect of high-fat feeding on circulating EVs, all cohorts in this study were pre-conditioned to handing (bleeding, scruffing, mock-injections etc.) for 2-3 weeks before they were placed on diet. We found that within a single day on a high-fat diet there was a surge of total EVs in the blood compared to chow fed mice (**Fig. 2E**). For both males and females, there was a compensatory normalization of circulating EVs at 7 days on diet (**Fig. 2E**). This effect was short-lived and plasma EVs were increased in the males on a high-fat diet by 3-fold at 2 weeks, peaking to 4-fold at 6 weeks, and maintaining an ∼2-fold increase through the 18-week diet (**Fig. 2F**). A similar result was observed in females (**Fig. 2F**). Plasma adipoEVs and adipoEV-mitos followed the same pattern as total EVs, increasing both acutely with high-fat feeding and remaining elevated above chow fed mice over most of the diet duration (**Fig. 2G-J**). Interestingly, total mitochondria in the blood did not change much over the course of the high-fat diet in males but showed a significant increase at 18 weeks (**Fig. 2K and L**). In females on a high-fat diet, total circulating mitochondria was consistently lower over the diet duration than those on chow (**Fig. 2K and L**). These data suggests that the levels of blood adipoEV-mitos and total mitochondria are differentially regulated. No consistent change in EVs from red blood cells, endothelial cells, or platelets were detected over the time course of high-fat feeding (**Fig. S3 A-C**).

**Figure 2.**
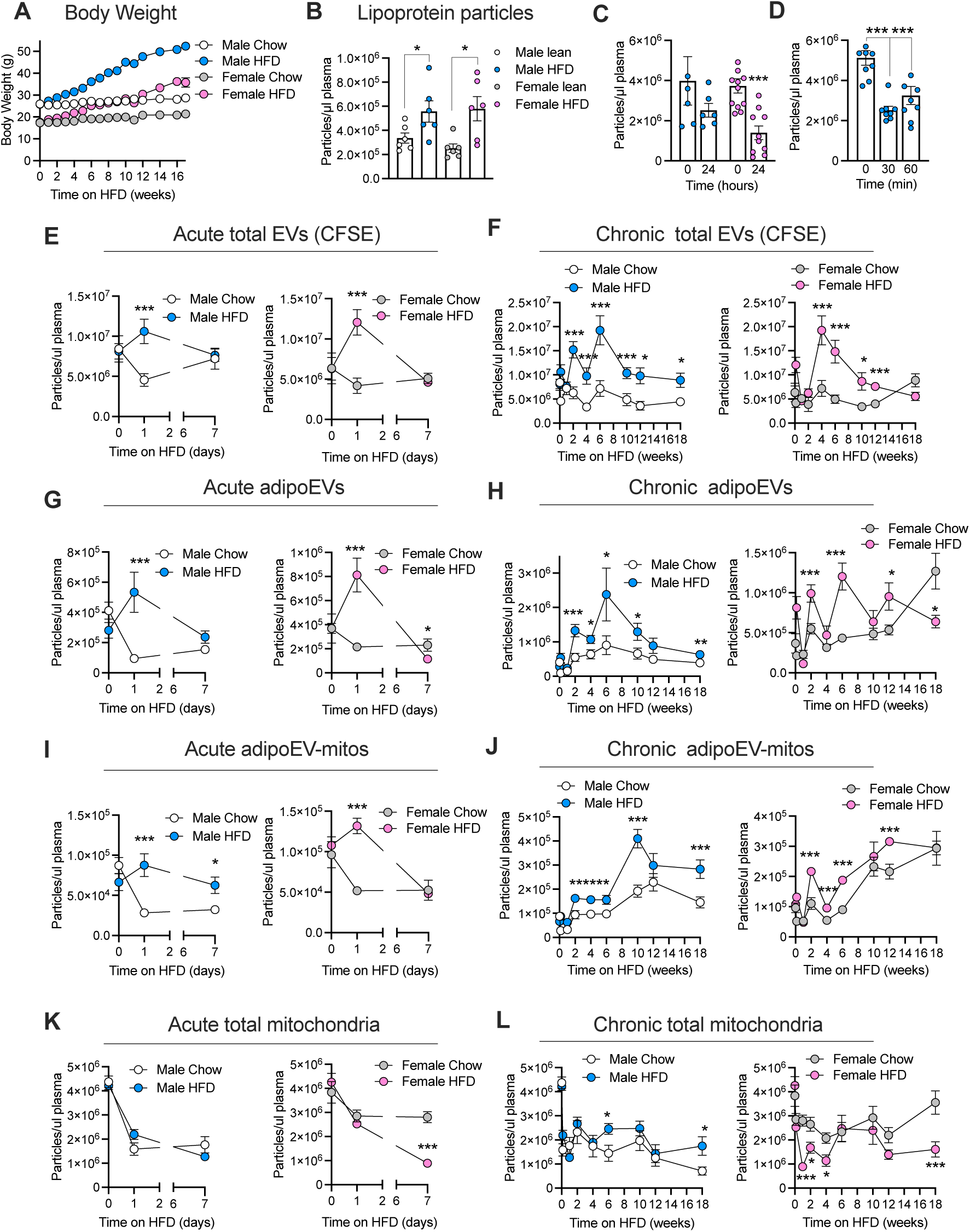
Circulating adipocyte EVs and EV-mitos are increased with high-fat feeding early and persistently. **A.** Body weights for a cohort of WT male and female (N=6 for each sex and group). **B.** Nano particle flow cytometry for lipoprotein markers in mice plasma (APOB/E antibodies). Mice were on diets for 6 weeks. **C-D** Total EVs in plasma (CFSE labeled) were quantified at day 0 and day 1 of the feeding experiment (**C**) or at baseline, and 30 or 60 minutes after bleeding (**D**). Plasma as collected from male and female cohorts (N=6) via tail vein bleed over the indicated time course after diet initiation (chow or high-fat diet; HFD). **E-L** EVs were analyzed for total EVs (CFSE-stained; **E-F**), adipocyte EVs (adipoEVs; APN/PLN1^+^; **G-H**), adipocyte EV-mitos (adipoEV-mitos; APN/PLN1^+^, VDAC^+^; **I-J**), and total mitochondria (VDAC^+^, **K-L**). **E, G, I** and **K** are acute timepoints. **F, H, J, L** are chronic timepoints. Data are presented as mean ± s.e.m. \**P* < 0.05, *** *P* < 0.001.

Because obesity etiology has a strong inflammatory component, we also quantified total immune cell EVs (CD45) and macrophage-specific EVs (F4/80) in mouse plasma. Macrophage EVs displayed the same increase as adipocyte EVs throughout the full diet duration (**Fig. 3A**). A small population of EVs that carried both the adipocyte markers and F4/80 was considered to be adipose tissue macrophage (ATM)-derived EVs. Circulating ATM EVs did not change in the first 6 weeks of high-fat feeding but robustly increased from 10-18 weeks on diet (**Fig. 3B**). Surprisingly, total immune cell EVs were unchanged throughout the high-fat diet timecourse (**Fig. 3C**). AdipoEVs, adipoEV-mitos, ATM-EVs, and total mitochondria are strongly correlated with fat mass (**Fig. 3D**).

**Figure 3.**
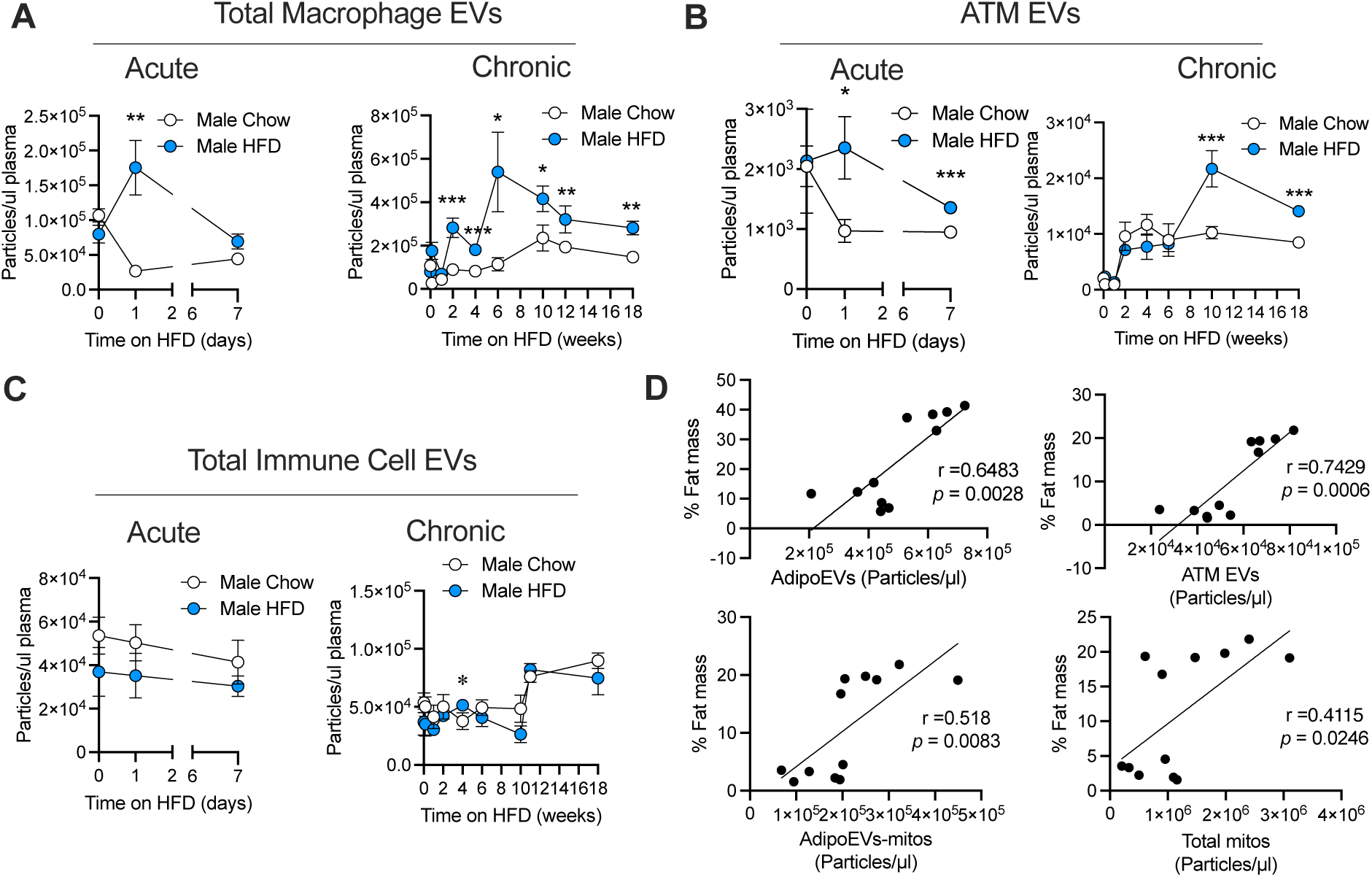
Macrophage EVs increase early and persistently in obesity. Plasma from mice on the chow (N=6) or high-fat diet (N=6) for the indicated times were analyzed for total macrophage EVs (**A**), adipose tissue macrophage EV (ATM EVs; **B**), or total immune cell EVs (**C**). **D**. correlation of % fat mass with adipoEV, ATM EV, adipoEV-mitos, or total mitochondria in plasma. Data are presented as mean ± s.e.m. \**P* < 0.05, ** *P* < 0.01, *** *P* < 0.001.

### AdipoEVs correlate with insulin resistance in people with obesity

To understand if similar changes occur in human obesity plasma samples from three groups of study participants were collected: metabolically healthy and lean (MHL; N=14), metabolically healthy obesity (MHO; N=14), metabolically unhealthy obesity (MUO; N=14). All groups were matched on age and sex (Table 1). The MHO and MUO groups were matched on body mass index (BMI) and percent body weight as fat and the MHL and MHO groups were matched on whole-body insulin sensitivity, assessed by the hyperinsulinemic-euglycemic-clamp procedure (Table 1). Circulating cell type-specific EVs were quantified as for mouse samples. Lipoproteins (APOB/E) increased in both MHO and MUO plasma compared to MHL (**Fig. 4A**). Both total EVs and adipoEVs were dramatically increased in the MUO group compared to MHL or MHO (**Fig 4B-C**). Like mice, lean human adipoEVs made up 10.7% ± 2.3% of total circulating EVs. AdipoEV-mitos trended towards an increase in the MHO group and were significantly higher in the MUO group compared to MHL (**Fig. 4D**). Likewise, total mitochondria were increased in the MUO group compared to other groups (**Fig. 4E**). Total immune cell EVs and adipose tissue immune cell EVs were significantly higher in MUO compared to MHO participants (**Fig. 4F-G**). Finally, endothelial cell EVs displayed a significant increase in MHO and an even greater increase in the MUO group compared to MHL (**Fig. 4H**). These data suggests that total EVs, adipocyte EVs, and immune cell EVs may be biomarkers that distinguish between healthy and unhealthy obesity in humans.

**Figure 4.**
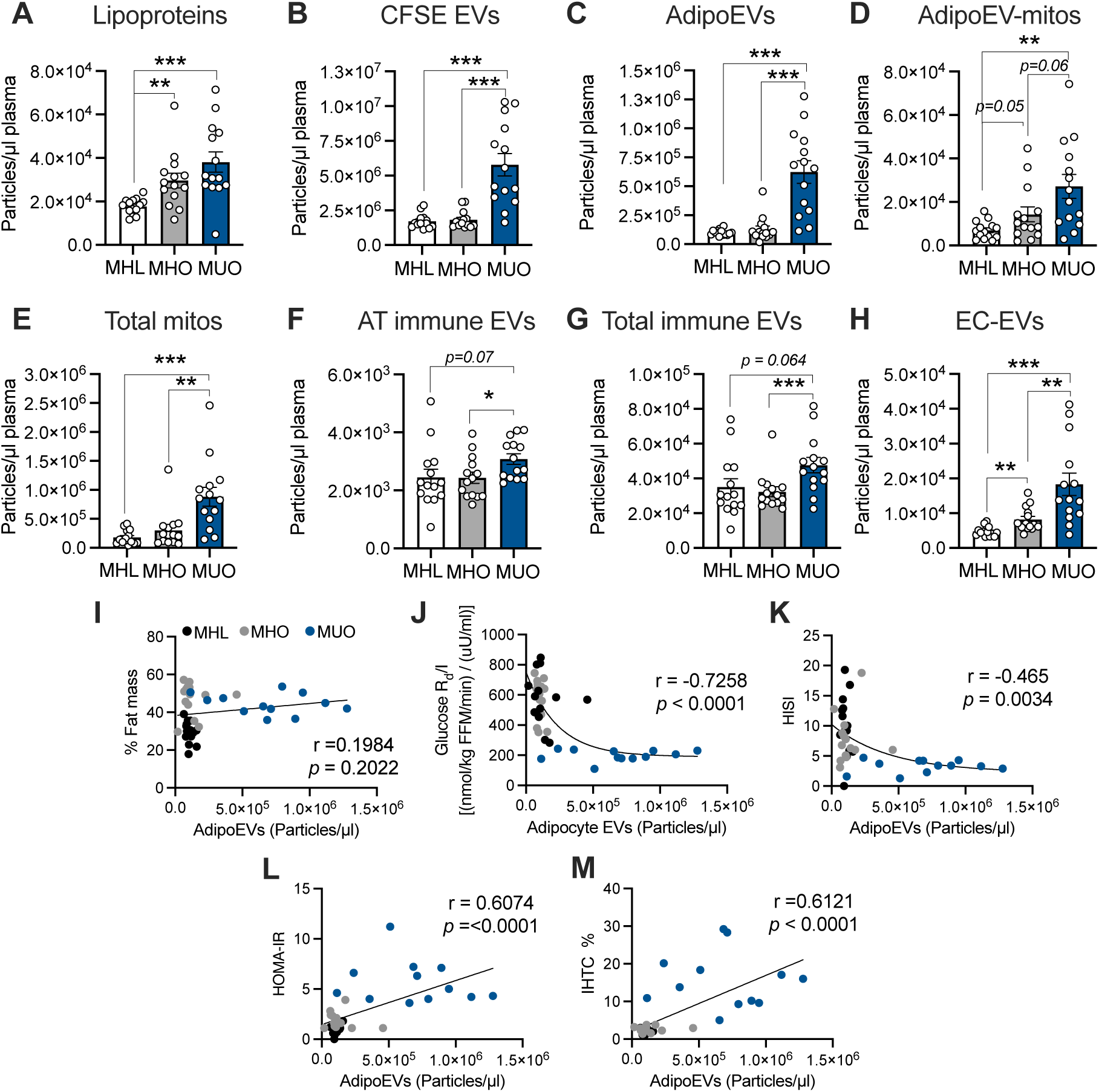
AdipoEVs correlate with insulin resistance in people with obesity. Plasma samples from metabolically healthy lean (MHL), metabolically healthy obese (MHO) and metabolically unhealthy obese (MUO) patients were analyzed by nano flow cytometry for **A.** Lipoproteins (APOE/B^+^), **B.** total EVs (CFSE^+^), **C.** adipoEVs (APN/PLN1^+^), **D.** adipoEV-mitos (APN/PLN1^+^, VDAC^+^), **E.** total mitochondria (VDAC^+^), **F.** Adipose tissue (AT) immune cell EVs (APN/PLN^+^, CD45^+^), **G.** total immune cell EVs (CD45^+^), or **H.** Endothelial cell (EC) EVs (CD31^+^/CD45^-^). AdipoEVs correlations with **I.** % fat mass **J.** insulin sensitivity, **K.** hepatic insulin sensitivity index (HISI), **L.** homeostatic model assessment of insulin resistance (HOMA-IR), **M.** Intrahepatic triglyceride (IHTC) content. Data are presented as mean ± s.e.m. \**P* < 0.05, ** *P* < 0.01, *** *P* < 0.001.

**Table 1.** Body composition and metabolic characteristics of the participants Data are expressed as mean ± SEM. FFM, fat-free mass. HISI, hepatic insulin sensitivity index. HOMA-IR, homeostasis model assessment of insulin resistance IHTG, intrahepatic triglyceride. Oral glucose tolerance test, OGTT. ^a^*P* ≤0.05 value significantly different from MHL value. ^b^*P* ≤0.05 value significantly different from MHO value

|  | MHL<br>(n=14) | MHO<br>(n=14) | MUO<br>(n=14) |
| --- | --- | --- | --- |
| Age (yr) | 36.7 ± 2.6 | 40.7 ± 2.0 | 40.6 ± 2.3 |
| Sex (Male/Female) | 4/10 | 4/10 | 4/10 |
| Body mass index (kg/m<sup>2</sup>) | 22.7 ± 0.4 | 36.5 ± 1.3<sup>a</sup> | 36.3 ± 1.1<sup>a</sup> |
| Body mass (kg) | 61.3 ± 1.7 | 106.5 ± 4.6<sup>a</sup> | 102.9 ± 3.2<sup>a</sup> |
| Fat-free mass (kg) | 42.7 ± 1.7 | 55.9 ± 3.4<sup>a</sup> | 55.7 ± 2.0<sup>a</sup> |
| Body fat (%) | 30 ± 1 | 47 ± 2<sup>a</sup> | 46 ± 2<sup>a</sup> |
| IHTG content (%) | 1.9 ± 0.2 | 2.7 ± 0.2 | 15.1 ± 2.0<sup>ab</sup> |
| HbA1c (%) | 5.0 ± 0.1 | 5.0 ± 0.1 | 5.5 ± 0.1<sup>ab</sup> |
| Plasma triglyceride (mg/dL) | 77 ± 8 | 63 ± 7 | 153 ± 12<sup>ab</sup> |
| Fasting glucose (mg/dL) | 85 ± 1 | 87 ± 1 | 97 ± 3<sup>ab</sup> |
| HOMA-IR | 1.2 ± 0.1 | 1.9 ± 0.2 | 5.6 ± 0.6<sup>ab</sup> |
| OGTT 2-h plasma glucose (mg/dL) | 99 ± 5 | 97 ± 4 | 171 ± 8<sup>ab</sup> |
| HISI:1,000/(μmol/kg FFM/min)*(μU/mL) | 10.6 ± 1.1 | 7.8 ± 1.1 | 3.2 ± 0.3<sup>ab</sup> |
| Whole-body insulin sensitivity:<br>(nmol/kg FFM/min) / (μU/mL) | 574 ± 36 | 572 ± 46 | 198 ± 9<sup>ab</sup> |

Unlike our mouse cohorts, human adipoEVs did not correlate with % fat mass (**Fig. 4I**). Instead adipoEV levels had a strong negative correlation with whole body insulin sensitivity (Glucose R_d_/I [(nmol/kg FFM/min) / (uU/ml)]) and the hepatocyte insulin sensitivity index (HISI; **Fig. 4J and K**). Consistent with this, adipoEVs had a strong positive association with the homeostatic model assessment of insulin resistance (HOMA-IR; **Fig. 4L**). AdipoEVs also were strongly correlated with percent intrahepatic triglyceride (IHTC) content (**Fig. 4M**). These data are all consistent with mouse studies that show injection with exogenous adipoEVs results in whole body insulin resistance, hepatocyte insulin resistance ^2,14–16^.

### Circulating EV levels are regulated by metabolic shifts

To begin to understand which physiological contexts endogenous adipoEVs may signal, we determined how plasma levels of adipoEVs change with major whole-body metabolic shifts like during feeding/fasting or diurnally. We fasted half of each cohort for 16 hours, followed by 3 hours of re-feeding their respective diets. The remaining mice were kept fed *ad libitum*. Blood was sampled from fasted, re-fed mice, and *ad libitum* fed mice sampled at the same time for each condition. Lean male and female mice displayed a striking reduction in total circulating EVs, adipoEVs, and adipoEV-mitos with fasting (**Fig. 5A-C**). The effect on total EVs was more dramatic in high-fat fed mice (**Fig. 5A**). EV levels were restored with re-feeding in lean mice (**Fig. 5A-B**). In high-fat diet mice, total EVs and adipoEVs were also restored but to a level significantly higher than *ad libitum* fed mice (**Fig. 5A-B**). Interestingly, total macrophage EVs were unaffected by fasting in both male and female lean mice (**Fig. 5D**). However, only obese female mice displayed a significant fasting-induced reduction in macrophage EVs (**Fig. 4D**). Refeeding triggered elevation of macrophage EVs levels above baseline in both obese males and females (**Fig. 5D**).

**Figure 5.**
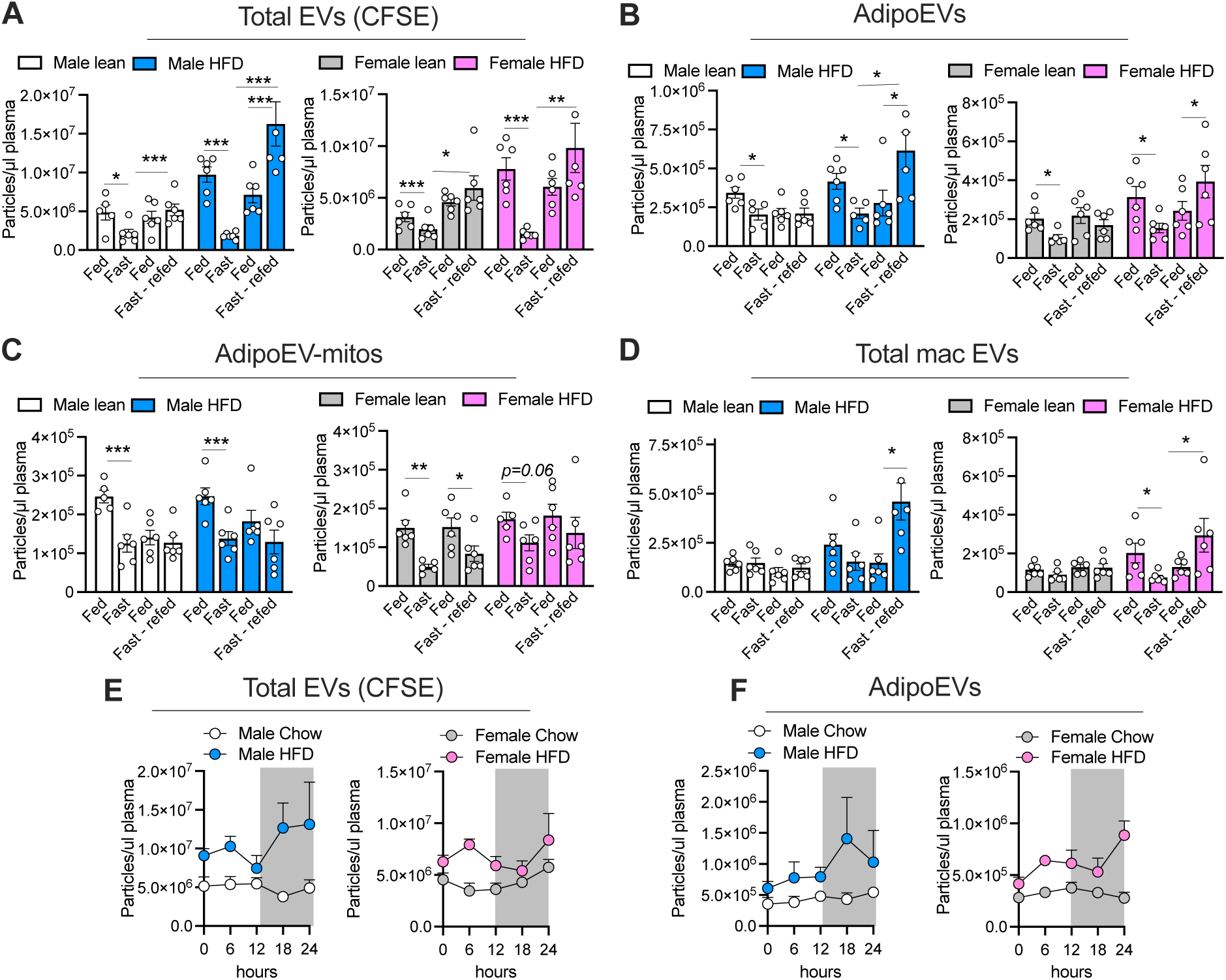
Circulating EVs and adipoEVs levels are regulated by the feeding/fasting transition. **A-D** Mice were on either a chow or high-fat diet for 7 weeks. Mice were either fed *ad libitum* their respective diets, or fasted for 16 hours then re-fed the indicated diet for 3 hours. Plasma samples in each metabolic state were analyzed for **A.** total EVs (CFSE^+^), **B.** adipoEVs (APN/PLN1^+^), **C.** adipoEV-mitos (APN/PLN1^+^, VDAC^+^), or **D.** macrophage (mac) EVs (F4/80^+^). **E** and **F.** Mice were on either chow of high fat diet for 9 weeks. Plasma was sampled every 6 hours over a 24-hour time course and analyzed for total EVs (**E**) or adipoEVs (**F**). Data are presented as mean ± s.e.m. \**P* < 0.05, ** *P* < 0.01, *** *P* < 0.001.

We next determined if adipoEVs are regulated diurnally. We sampled blood from lean or obese, male or female mice, every 6 hours over a 24-hour period. We did not detect dramatic circadian changes in total EVs or adipocyte-specific EVs in lean mice, although total EVs were at the lowest at the 18-hour timepoint (12 am) in males and at the light-to-dark transition point (6 pm) in females (**Fig. 5E**). AdipoEVs did not display circadian regulation in lean males but tended to peak at the light-to-dark transition (6 pm) in lean females. In obese mice, total EVs and adipoEVs displayed completely altered circadian patterns in both obese males and females compared to lean mice (**Fig. 5E-F**).

### Macrophage-mediated EV clearance regulates plasma levels of EVs

We wanted to determine how circulating EV levels are regulated. AdipoEV levels in mouse plasma highly correlate with total plasma EVs (**Fig. 6A**). This suggests that there may be a universal mechanism of EV clearance that also regulates adipoEV plasma levels. Several studies have shown that exogenous EVs generated from cell lines, injected i.v. into mice are cleared from circulation strikingly fast, and that macrophages are likely responsible^33^. In addition, depletion of macrophages leads to a marked increase in the release of adipocyte-derived mitochondria into blood^34^. To test if this is true for endogenously produced “self” EVs, we depleted macrophages and monocytes by treating mice with clodronate-loaded liposomes. Male mice fed a chow or high-fat diet for 6 weeks were injected with clodronate every 3 days to maintain macrophage depletion (**Fig. 6B**). Macrophage depletion resulted in a dramatic increase in total circulating EVs and adipoEVs, an effect that was augmented in high-fat fed mice (**Fig. 6C-D**). Total EVs in high-fat diet-fed, clodronate treated mice peaked at levels that were 10-fold higher than that seen in high-fat diet mice injected with the liposome controls at day 2 of treatment (**Fig. 6C**). Hepatocyte and endothelial cell-derived EVs also increased substantially in circulation at day 2 of clodronate treatment in mice on chow and more so on high-fat diet (**Fig. 6E** and **F**) This effect was not seen with myocyte EVs (**Fig. 6G**). Interestingly, clodronate did not reduce the number of macrophages in the gastrocnemius muscle, but did deplete macrophages in both the liver and adipose (**Fig. 6P and Fig. S4**). Clodronate caused the most dramatic increase in blood cell (platelets and red blood cells) EVs (**Fig. 6H-I**). No effect of clodronate was detected on lipoprotein particles (**Fig. 6J**). These data suggest that either more EVs are being produced, or they are cleared slower when macrophages are absent. To measure the clearance rate of EVs in lean and overweight mice, we fluorescently labeled total EVs in mouse plasma with CFSE, purified EVs away from free dye with SEC, and retro-orbitally injected the labeled EVs into lean or overweight mice +/- clodronate. We collected tail blood at the indicated times and quantified the number of remaining CFSE^+^ EVs by nano-flow cytometry. Surprisingly, within 10 minutes, over 90% of injected EVs were cleared from circulation in lean mice (**Fig. 6K**). Mice fed a high-fat diet displayed a significantly higher level of residual CFSE^+^ EVs suggesting a slower clearance rate compared to lean mice (**Fig. 6K**). Depletion of macrophages in lean mice resulted in reduced EV clearance rate (**Fig. 6L**). Interestingly, high-fat diet fed mice displayed the same initial rate of EV clearance in the presence or absence of macrophages (**Fig. 6M**), suggesting EV clearance may be less reliant on macrophages in mice on a high-fat diet. As further evidence of this, in lean mice, the fasting-induced reduction in total circulating EVs is absent in clodronate-treated mice, which instead displayed a slight increase in total EVs with fasting (**Fig. 6N**). In contrast high-fat diet fed mice treated with clodronate maintained fasting-associated reduction in total EVs, although to a lesser degree than that of liposome treated control mice (**Fig. 6N**). This effect was more dramatic for adipocyte-derived EVs, which did not change with fasting in clodronate-treated lean mice but were dramatically reduced in clodronate-treated high-fat diet-fed mice (**Fig. 6O**). Together these data suggest macrophages may be a key determinant of the circulating total EVs and adipocyte-specific EV levels in the lean state. With high fat feeding macrophage-mediated uptake does not contribute as much to EV clearance rate (**Fig. 6M**). Clodronate depletes both monocytes and macrophages. Macrophages were successfully depleted in the epididymal white adipose tissue (eWAT), but there was a compensatory increase in circulating monocytes (**Fig. 6P** and **Q**). This suggests that blood monocytes were likely not a major contributor to EV clearance rate from circulation. Instead, tissue macrophages are playing a prominent role.

**Figure 6.**
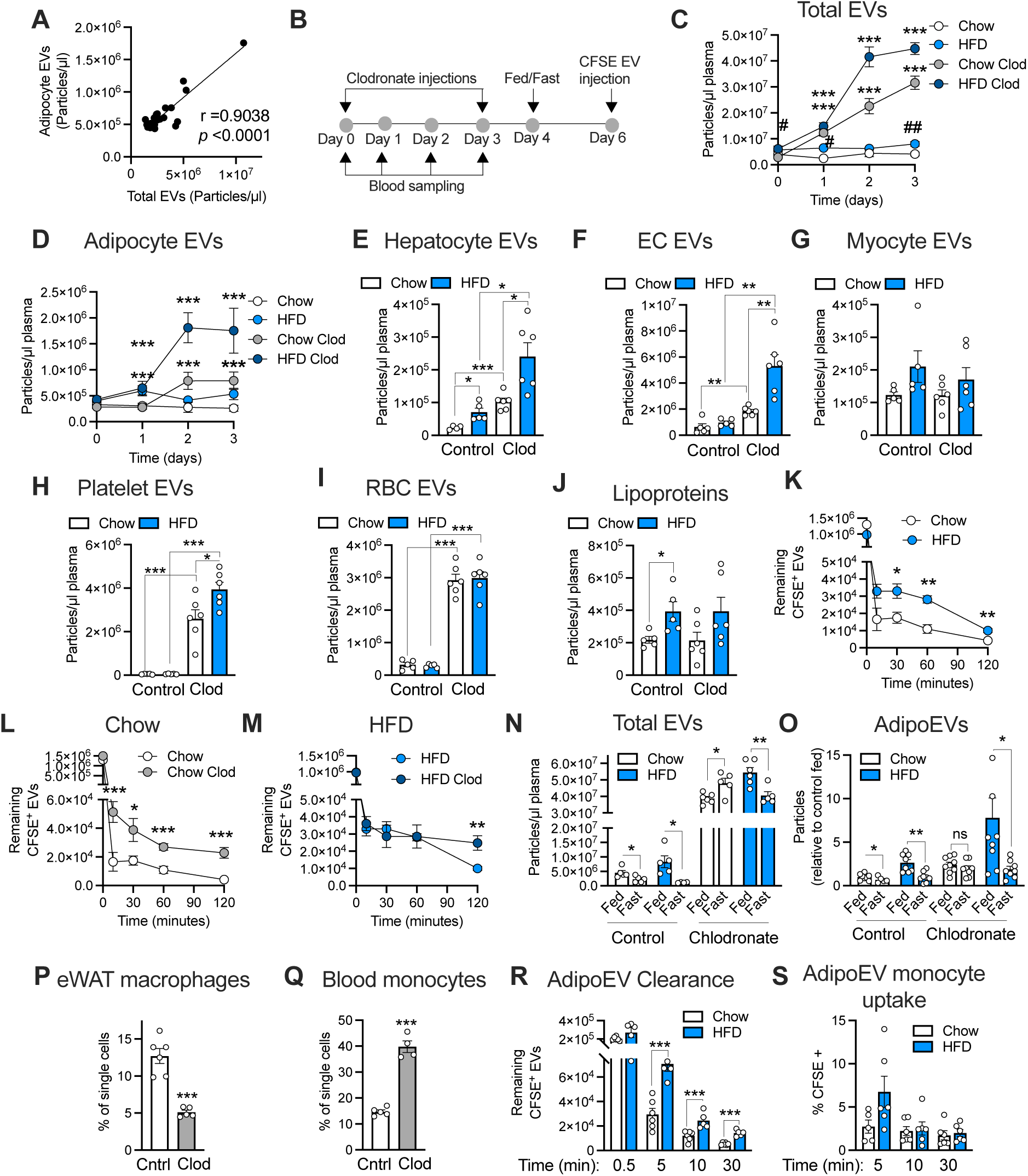
Macrophage-mediated clearance of EVs is reduced with high-fat feeding. **A.** AdipoEVs (APN/PLN1^+^) correlate with total EVs (CFSE^+^). **B.** Timeline of clodronate experiment. Mice were on a chow of high fat diet for 6 weeks at which point 5µl/gram mouse body weight of either control liposomes or clodronate liposomes were injected into mice at day 0 and day 3. Blood was sampled at the indicated timepoints. At day 4 of macrophage depletions and fed/fasted study was conducted. At day 6, mice were injected i.v. with purified plasma or adipocyte EVs labeled with CFSE for determination of EV clearance rates. Total EVs (CFSE^+^; **C**) and adipoEVs (**D**) were quantified at the indicated timepoints. At day 2 post clodronate injection plasma samples were analyzed for **E.** hepatocyte EVs (ASGR1^+^/CD45^-^), **F.** endothelial cell (EC) EVs (CD31^+^/CD45^-^), **G.** myocyte EVs (MHC^+^, CD45^+^). **H.** platelet EVs (CD41^+^), **I.** red blood cell (RBC) EVs (TER119^+^), or **J**. lipoproteins (APOE/B^+^). **K-M** Mouse plasma EVs were harvested, labeled with CFSE, purified by SEC and injected retro-orbitally into mice on a chow or high fat diet (HFD) either treated with empty liposomes (Control) or clodronate liposomes (Clod). Tail blood was sampled at the indicated timepoints following CFSE^+^ EV injection and the CFSE^+^ EV remaining in the blood was quantified. **N-O** Mice treated with empty liposomes (Control) or clodronate liposomes were either *ad libitum* fed or fasted for 16 hours and refed for 3 hours. Plasma was sampled under each metabolic state and total EVs (CFSE; **N**) and adipoEVs (**O**) were quantified. At day 7 of the experiments (4 days after the last clodronate injection) mice were euthanized. **P.** Macrophages in the eWAT and **Q.** monocytes in the blood were quantified by flow cytometry in mice treated with control liposomes (Cntrl) or clodronate liposomes (Clod). **R.** EV clearance assay for adipocyte-derived CFSE^+^ EVs, as described in K-M. **S.** Blood monocyte uptake of adipocyte-derived CFSE^+^ EVs at the indicated timepoints after EV injection by flow cytometry. Data are presented as mean ± s.e.m. \**P* < 0.05, ** *P* < 0.01, *** *P* < 0.001.

To determine if adipocyte-specific EV clearance rates are slowed with high-fat diet, like total EVs (**Fig. 6K**), we labeled EVs purified from adipocyte culture media with CFSE and injected them retro-orbitally into mice. CFSE^+^ EVs were cleared from circulation remarkably fast in lean mice but notably slowed in mice on a high-fat diet (**Fig. 6R**). Blood monocytes took up these adipocyte EVs (CFSE^+^ monocytes), however, there was no differential uptake between lean and overweight mice (**Fig. 6S**). This is further evidence that monocyte uptake rates do not account for changes in clearance rate of adipoEVs between lean and high-fat fed mice.

To understand how tissue macrophage uptake of adipoEVs is altered in obesity we quantified the amount of adipocyte EVs taken up by adipose tissue immune cells. We used a mouse model with a doxycycline-inducible plasma membrane-targeted halo tag transgene (TRE-Halo) driven by adiponectin-rtTA in the presence of doxycycline. This adipocyte-specific halo tag incorporates well into adipocyte EVs as shown with immunogold stained electron micrograph of the sWAT extracellular space (**Fig. 7A**). We have previously used this mouse to track endogenously produced adipocyte EVs (halo^+^) to other tissue cells^35^ (**Fig. 7B**). Immune cells made up the greatest proportion of cells in eWAT, sWAT, and brown adipose tissue (BAT), that were positive for halo (received adipocyte EVs; **Fig. 7C**). Macrophages, dendritic cells, and neutrophiles displayed the highest level of adipocyte EV uptake among tissue cells (**Fig. 7C**). Depending on the adipose depot, 40-80% of all macrophages took up adipocyte EVs (**Fig. D-G**). 10-40% of all dendritic cells and neutrophiles took up adipocyte EVs (**Fig. 7E-G**). In obese mice, macrophage uptake of adipoEVs was reduced in both eWAT and BAT but not sWAT (**Fig. 7E-G**). Dendritic cell uptake was reduced with obesity only in eWAT (**Fig. 7E**). Lastly, neutrophil uptake of adipoEVs was suppressed in all adipose depots (**Fig. 7E-G**). Therefore, it is likely that several tissue resident immune cells, not just macrophages, are important for scavenging both blood-borne EVs but also newly produced EVs in the tissue. These immune cells take up less adipoEVs despite increased EV production from obese adipose tissue *ex vivo* (**Fig. 7H**). If we isolate adipose tissue EVs (ATEVs) directly from the adipose tissue, although there are more EVs in the entire fat pad with obesity, per mg tissue, there is either less (sWAT) or no difference (eWAT) in ATEVs in obese mice compared to lean (**Fig. 7I-J**). Therefore, even though obese adipose tissue has a higher rate of EV production the tissue retains less or the same amount of EVs, and displays reduced immune cell-mediated clearance. This suggests EVs are exiting the tissue, resulting in higher circulating levels. These data suggests that tissue immune cells may be a major regulator of inter-organ signaling of EVs by dictating the amount of EVs available to signal.

**Figure 7.**
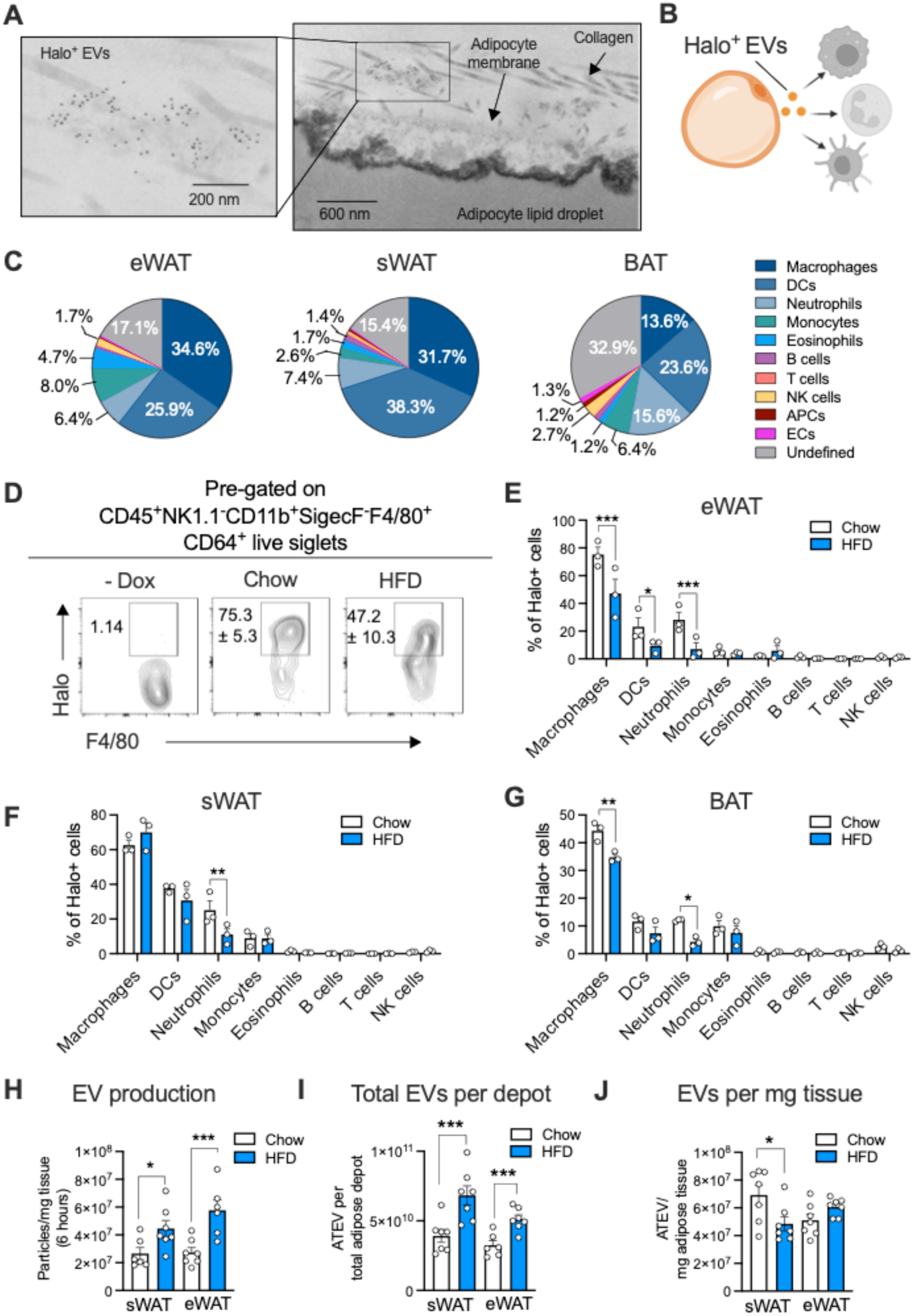
EV uptake by adipose tissue resident immune cells is reduced in obesity. **A.** A representative electron micrograph of the sWAT of adipocyte-specific, membrane-targeted halo tag transgenic mice. The Halo tag was immune-gold labeled. **B.** An illustration of the experimental premise, the halo tag is transferred from adipocytes in EVs to other tissue cells. **C.** Pie charts depicting the proportion of each cell type out of the total halo positive cells in the indicated tissue. (**D-G**) Membrane Halo male mice were fed with a chow or a high-fat diet (HFD) both with doxycycline (Dox) for 20 weeks. Mice that didn’t receive Dox served as negative control (-Dox). **D.** Representative flow plots showing the percentage of macrophages that captured the halo tag in eWAT. **E-G** The amount of the halo tag captured by the indicated immune cell types in eWAT (**E**), sWAT (**F**), and BAT (**G**). **H.** sWAT or eWAT explants were incubated for 6 hours in culture media and EVs were quanified by NTA. **I-J.** Adipose tissue EVs (ATEVs) were isolated from the sWAT or eWAT. EV count is displayed per entire adipose depot (I) or per mg adipose tissue (J). Data are presented as mean ± s.e.m. \**P* < 0.05, ** *P* < 0.01, *** *P* < 0.001. DCs, dendritic cells; NK cells, natural killer cells; APCs, adipocyte progenitor cells; ECs, endothelial cells.

## DISCUSSION

This study provides evidence that macrophages clear EVs released by local tissue cells (Fig. 7), or EVs coming to the tissue from circulation (Fig. 6K-M). Our data suggests that dendritic cells and neutrophiles likely play a similar role in scavenging EVs. Immune cells in general win in the competition for EV uptake within the tissue (Fig. 7C). The consequence is that, by intercepting EVs before they can interact with would-be target cells, immune cells are effectively a barrier to EV signaling. With high-fat feeding, macrophages in the eWAT and BAT take up less adipocyte EVs (Fig. 7), which, in addition to increased adipose tissue EV production, may allow adipoEVs to exit the tissue, accumulate in circulation, and enter other organs. This implies that adipoEV signaling may be restrained in a healthy state, but in a diseased tissue, as in obesity, macrophage function is altered accelerating EV signaling in the tissue and between organs. Although future work will have to determine if this model is true across all tissues and diseases, it is interesting that circulating EVs carry robust biomarkers of various tissue-specific diseases which are not, or lowly, present in healthy patient plasma EVs^36^. It is possible that disease-related immune cell dysfunction may facilitate the escape of EVs from the affected organ and appearance in circulation. This regulation at the level of the tissue microenvironment may also explain why we detect adipocyte and myocyte EVs at similar levels in the blood when muscle is reported to produce EVs at a faster rate than adipose^37^.

Macrophage-mediated regulation of EV levels seems to apply to most, if not all, EVs indiscriminate of origin cell (Fig. 6C-I). This may indicate a universal mechanism of EVs uptake by macrophages. Resident macrophages have been shown to take up adipocyte-derived, free extracellular mitochondria (not associated with an EV membrane) by a heparan sulfate (HS)-dependent mechanism^38^. Glycans, particularly HS, have also been shown to facilitate binding of EVs to recipient cells for endocytosis^39^. Furthermore, Borcherding et. al. reported that dietary long chain fatty acids inhibit adipose tissue resident macrophage uptake of extracellular, free mitochondria, allowing adipocyte mitochondria to enter the blood^34^. This inhibitory effect of fatty acids was dependent on HS. Because HS can bind to a wide range of proteins, it is possible that EV and EV-mito uptake by macrophages is also mediated by HS as a general pathway to clear extracellular material. If, like extracellular mitochondria, long chain fatty acids are responsible for the reduction in EV clearance we observe with high-fat feeding is yet to be determined. This HS-mediated mechanism of EV uptake implies macrophages are non-specifically scavenging EVs, but this does not rule out potential signaling effects of the EVs in immune cells. In fact, we know adipocyte EVs can change macrophage function^11,27^. Future work must determine the fate of adipocyte EVs once taken up by macrophages and if that changes in different metabolic conditions.

In recent years a growing number of studies have reported EV-mediated signaling for metabolic regulation. However, until recently, the inability to quantify cell type specific EVs below 200nm in a complex biofluid like plasma has restrained investigation of endogenously produced EVs. In this study, we used new flow cytometry detection technology to quantify almost the full size-range of cell-type specific EVs in plasma (80nm – 2µm). We found that adipocyte EVs and EV-mitos are dynamically regulated by dietary fat and feeding/fasting transitions. Us and others have shown increased adipocyte markers and adipocyte-derived mitochondria in the circulation with obesity^18,34,40^. We find here that adipocyte EVs, adipocyte EV-mitos, and macrophage EVs increase within 1 day of high-fat feeding and remain persistently elevated above the levels of lean mice during obesity progression. This suggests that adipocyte and macrophage EVs may participate in inter-organ signaling under physiological changes in dietary composition, not just in pathophysiological conditions. The nature of this early signaling is the topic of future studies. In human obesity, we found adipocyte EVs correlate positively with insulin resistance measures and liver lipid and negatively correlate with measures of insulin sensitivity (Fig. 4). Interestingly, human adipocyte EVs did not correlate with body fat mass as it did in mice. The human data suggests that it the number of adipocyte EVs in the blood is not a function of the mass of adipose tissue, but instead of metabolic dysfunction, potentially dysfunction at the level of adipose tissue. In the inbred mouse strain used in this study (C57B/6J), there are no healthy obese mice, therefore, adipose mass and metabolic health as determinants of circulating adipocyte EVs cannot be separated. Although we can’t conclude a causative role of adipocyte EVs in insulin resistance from our human dataset, there have been several studies in pre-clinical models and *in vitro* studies with human cells that have demonstrated role of adipocyte or adipose EVs in propagating insulin resistance and pro-inflammatory signals to other cells or organs^14–16,26,27,41^.

These pro-inflammatory, and insulin resistant signals from adipocytes EVs have been demonstrated with *in vivo* injections, which have shown surprisingly strong metabolic effects. However, there are concerns in the field that artifacts are created by injecting a supraphysiological dose of cell-type specific EVs. This is particularly important because the majority of what we know about EV function *in vivo* comes from injections of cell-type-specific EVs generated in culture, without any knowledge about what a physiologically relevant dose is. Because of this, chosen doses are wildly different between studies^42^. Therefore, a goal of this study was to provide an estimate of metabolic cell-type EVs in circulation of mice as a guide for future research (Table S1). This is the most conservative estimation, as it is likely that not every cell-specific EV will have a cell-type specific maker on/in it or have low levels of that maker that are not bright enough for us to detect. In addition, how well we detect cell makers is dependent on antibodies that have different binding affinities and fluorophores for each marker, which can affect how many positive EVs we detect. We estimate that adipocyte EVs make about 10% of total EVs (CFSE positive particles) in mouse and human plasma. Many have speculated that adipocytes contribute the majority of small EVs (<200nm diameter) to circulation based on a study by Thomou et. al. that demonstrated at least 50% of small EV-associated micro RNAs (miRNAs) originate from adipocytes^8^. This may reflect the high enrichment of miRNAs in adipocyte EVs compared to other cell-type-specific EVs but is not necessarily predictive of adipocyte EV number. Additionally, miRNAs are concentrated in the smallest EVs, which our flow cytometry analysis does not capture. In contrast, studies using computational approaches that sequence small EV RNAs from human plasma as predictors of cell origin suggest most plasma EVs are derived from hematopoietic cells, which we also found in this study, but only 0.2-1% from solid organs^19,43^. Although Li et. al. report that ∼80% of those solid tissue EVs are derived from adipose tissue, this is still substantially less than we estimate. One reason for this is these studies, are using bulk RNA sequencing data from total isolated EVs. This is an excellent approach to identify cell of origin but cannot distinguish between there being a higher number of a cell-type specific EVs or more copies of that specific marker in each EV. Furthermore, sample processing can also affect the estimation because the lipid content of adipocyte EVs causes them to float during ultracentrifugation^11^ a method of EV isolation used in these computational studies. Affinity-based EV purification was also employed, which too only recovers a sup-population of EVs. Therefore, the combination of our single EV detection approach and SEC-mediated EV isolation allowed us to detect a population of EVs that may be routinely discarded in other studies.

EVs have been described as “the next frontier in endocrinology”^44^. Just like the development of the radioimmunoassays in the 1950s allowed scientists to uncover the function of endocrine hormones, we expect the ability to quantify circulating EVs from metabolically important cells across physiological conditions will advance our understanding of endogenous EV signaling in health and metabolic disease

## METHODS

### Animals

All procedures performed on animals were approved by the Institutional Animal Care and Use Committee (IACUC) of Washington University in St. Louis School of Medicine. Cohorts of C57BL/6J mice used for plasma EV determination were acquired from Jax Laboratories (Strain # 000664). Cohorts were received at 4 weeks of age and allowed to acclimate to the facility for 1 week. Mice were then conditioned to handling stress 3-4 times a week through mock bleeding and/or scruffing and saline injections in line with the intended future procedures. Mice were started on a high fat diet (60% fat; Bio-Serv S1850) or matched control diet (Bio-Serv S4031) at 8 weeks of age. Adipocyte-specific PPARγ knockout mice were generated by crossing PPARγ floxed mice (Jax Laboratories; 004584) with adiponectin Cre mice (Jax Laboratories; 028020). For EV tracing the adiponectin-rtTA mouse was crossed with the TRE-membrane halo mouse for a doxycycline-inducible model of adipocyte labeling as previously described^35^.

### Clodronate depletion of macrophages

Mice were maintained on a chow or high fat diet for 6 weeks. Macrophage and monocyte depletion was initiated by retro-orbital injection of 5 µl/gram body weight clodronate liposomes or control, empty liposomes (FormuMax Scientific, F70101C-NC-10). Mice received a second injection intraperitoneally 10 hours later and a 3^rd^ injection 3 days later.

#### Body composition analysis

Mice were weighed, and body composition was determined by EchoMRI-100H 2n1 with a horizontal probe configuration (EchoMRI, Houston, TX).

### Human studies

A total of 42 men and women participated in this study, which was approved by the Human Research Protection Office at Washington University School of Medicine (WUSM) in St. Louis, MO and registered in ClinicalTrials.gov (NCT02706262). All subjects provided written consent, informed consent before their participation. A comprehensive medical evaluation was completed in the Clinical Translational Research Unit (CTRU) at WUSM by all subjects after an ∼12-h overnight fast to determine eligibility, which included a history and physical examination, standard blood tests, a 2-h oral glucose tolerance test (OGTT), and magnetic resonance imaging (MRI) (3-T MAGNETOM Vida; Siemens) to determine intrahepatic triglyceride (IHTG) content. During the screening visit, blood samples were also collected to analyze EDTA-plasma EVs and to determine plasma glucose and insulin concentrations, which were used to determine homeostasis model assessment of insulin resistance (HOMA-IR) by dividing the product of the fasting plasma insulin (in µU/mL) and glucose (in mmol/L) by 22.5^45^. Three groups of participants separated by adiposity and metabolic health were enrolled: 1) metabolically healthy lean (MHL; n=14), defined as a body mass index (BMI) 18.5-24.9 kg/m^2^; IHTG content <5%, plasma triglyceride <150 mg/dL, fasting plasma glucose < 100 mg/dL, 2-h OGTT plasma glucose <140 mg/dL, and hemaglobin A1c (HbA1c) ≤5.6%; 2) metabolically healthy obese (MHO; n=14), defined as BMI 30-49.9 kg/m^2^; IHTG content <5%, plasma triglyceride <150 mg/dL, fasting plasma glucose <100 mg/dL, 2-h OGTT plasma glucose <140 mg/dL, and HbA1c ≤5.6%; and 3) metabolically unhealthy obese (MUO; n=14), defined as BMI 30-49.9 kg/m^2^, IHTG content ≥5% and HbA1c ≥5.7% or fasting plasma glucose ≥100 mg/dL or 2-h OGTT plasma glucose ≥140 mg/dL. No subject had a history of diabetes, liver disease other than MASLD, took medications known to affect the study outcomes, or consumed excessive amounts of alcohol (>21 units of alcohol/week for men and >14 units for women).

### Body composition and insulin sensitivity assessments

Body fat mass and fat-free mass (FFM) were determined using dual-energy X-ray absorptiometry (Lunar iDXA, GE Healthcare). A hyperinsulinemic-euglycemic clamp procedure was used to assess hepatic and whole-body insulin sensitivity. Participants were given standard meals at 1900 h after admission to the CTRU on day 0 and at 0700, 1300, and 1900 h on day 1. At 0715 h on day 2, a primed (8 µmol/kg body mass), continuous (0.08 µmol/kg body mass/min) infusion of [U-^13^C]glucose was started. At 210 min after the start of the glucose tracer infusion (end of basal period), insulin was infused at a rate of 50 mU/m^2^ body surface area (BSA) initiated with a two-step priming dose: 200 mU/m^2^ BSA/min for 5 min followed by 100 mU/m^2^ BSA/min for 5 min. Dextrose (20%), enriched to ∼1% with [U-^13^C]glucose to minimize changes in plasma glucose isotopic enrichment, was infused at a variable rate to maintain plasma glucose concentration at ∼100 mg/dL during insulin infusion (for 210 min). Blood samples were obtained before starting the glucose tracer infusion and every 6 to 7 min during the last 20 min of the basal and insulin infusion stages to determine plasma insulin concentrations and glucose tracer-to-tracee ratios (TTR) by using gas-chromatography/mass spectrometry (GC-MS; Hewlett-Packard MSD 5973 system with capillary column) as previously described^46^. The hepatic insulin sensitivity index (HISI) was calculated as the inverse of the product of plasma insulin concentration and the endogenous glucose rate of appearance (R_a_) into the systemic circulation, determined by dividing the glucose tracer infusion rate by the average plasma glucose TTR during the last 20 min of the basal period of the HECP^47^. Total glucose rate of disappearance (R_d_) during insulin infusion was assumed to be equal to the sum of endogenous glucose rate of appearance into the bloodstream and the rate of infused glucose during the last 20 min of the clamp procedure. Skeletal muscle insulin sensitivity was calculated as glucose R_d_ expressed per kg FFM divided by the average plasma insulin concentration (glucose R_d_/I) during the final 20 min of the clamp procedure^47^.

### Preparation of 96 well Size Exclusion column plate

Sepharose® CL-6B (Sigma; CL6B200) resin was de-gassed under a vacuum and allowed to sit at 4°C overnight. Storage solution was removed, and 0.2 µm-filtered water was added 1:1 to make a 50% slurry. A 96 well filter plate (Agilent; 201718-100) was washed by adding 600 µl of 50% isopropanol to each well and allowed to fully drain. The CL-6B slurry was then inverted gently to suspend the resin and 850 µl was transferred to each well of the filter plate. The liquid was allowed to drain from the filter plate leaving 500 µl packed resin. The resin was washed one time with 600 µl 50% isopropanol, followed by two times with 0.2 µm-filtered water, and finally, five times with DPBS (Sigma). Before samples were added to the wells, the resin was washed once more with 600 µl of 0.02 µm-filtered DPBS.

### Sample processing for flow cytometry

Mouse blood was collected from the tail into EDTA tubes (STARDENT) that were pre-loaded with 4 µl heparin (1000 IU/ml stock solution). Mice were always bled at the same time of day (9:00 am) unless indicated differently.

Bleeds performed during the dark cycle were done under red light. Blood was placed on ice immediately and plasma was separated as soon as all the samples were collected for a given timepoint. Blood was centrifuged at 2,500 xg once at 4°C for 10 minutes. ∼20 µl plasma was recovered for each sample. Care was taken to recover the floating lipid layer and not disrupt the cell pellet. Plasma was aliquoted and stored in −80°C. No sample was freeze/thawed more than twice for flow cytometry. For carboxyfluorescein succinimidyl ester (CFSE) staining, an aliquot was defrosted and 2 µl of each sample was transferred into an 8-tube strip containing 48 µl of 25 µM CFSE (Thermo) diluted in 0.02 µm-filtered DPBS. Samples were incubated at 37°C for 30 minutes and each was loaded onto a SEC column to remove excess dye. Up to 96 samples were loaded at one time by multichannel pipette and allowed to soak into the resin. Once the columns stopped dripping, the filter plate was transferred to a collection plate and 300 µl 0.02 µm-filtered DPBS was added to each well and allowed to drain into the collection plate. The eluted EVs were either directly measured by flow cytometry or diluted before measurement (1:2 – 1:5 in 0.02 µm-filtered DPBS). Reported values were corrected for dilution factor. For antibody labeling, 2 µl plasma was combined with 23 µl Fc block (1:200) diluted in 0.02 µm-filtered DPBS. For the adipocyte, hepatocyte, or myocyte antibodies, 0.00002% triton X-100 was added to permeabilize EVs for detection of luminal markers. Plasma was incubated in this initial solution for 5 minutes then mixed with 25 µl of a 2X antibody solution. The 2X antibody solution was made by combining antibodies in each panel together at a high concentration (20X) with brilliant stain buffer (BD Biosciences) for 10 minutes. Following this incubation the antibodies were diluted in 0.02 µm-filtered DPBS 10X so that the concentration of each antibody is 2 times the desired final concentration. Once the antibody solution was mixed with samples, they were incubated overnight at 4°C in the dark, rocking. Before flow cytometry analysis, EVs were purified away from free antibody and plasma proteins with the SEC columns as described for the CFSE stain. See supplementary Table S3 for antibodies used and concentrations for each panel. To establish the panels, antibodies were titrated with 2 µl of plasma avoid saturation of the signal. The adiponectin antibody was conjugated to CL647 (Biotium mix-n-stain antibody labeling kit). The APOB and APOE antibodies were labeled with CoraLite Plus 750 (Proteintech FlexAble antibody labeling kit) fresh before each experiment.

Adipose tissue mitochondria were labeled with antibodies and processed as described above for EVs. Antibodies were titrated with ∼6×10^10^ mitochondria (as measured by nanoparticle tracking analysis). The final dilution of mitochondria for flow cytometry analysis after SEC was 1:150.

### EV flow cytometry

Flow cytometry analysis was conducted with the Cytek Northern Lights spectral flow cytometer with enhanced small particle detection (ESP) with fluorescence gain setting set to 4 times the default setting (CytekAssaySetting). Parameters were optimized using Flow Cytometry Sub-micron Particle Size Reference Kit (ThermoFIsher). SEC-purified EVs from mouse plasma were titrated to determine the appropriate dilution to prevent swarming (abort rate remained under 10% of the event rate). The violet side scatter laser was used as the trigger channel and the threshold was set to 600. The side scatter trigger channel was calibrated for EV size determination with Rosetta Calibration beads (Exometry) using the recommended parameters (EV model/refractive index; Mie modeling). See Table S2 for MIFlowCyt-EV check list^48^. Control experiments were conducted including, buffer only, reagent only, isotype controls (Figure S1-2). Fluorescence minus one (FMO) controls were used to confirm gating and avoid artifacts due to fluorophore interactions. Finally, the same mouse plasma sample was run with each batch to assess day to day variability and inter-data set comparability.

### Cell Flow Cytometry

Flow cytometry was conducted from cells from whole blood and from stromal vascular fraction from adipose depots. Whole blood was collected from mice by tail bleed in EDTA tubes. Approximately 10 µL of blood was drawn from each mouse, mixed, and immediately placed in ice till further processing. Red blood cells were lysed with 1:50 red blood cell lysis buffer (Roche, 11814389001) and incubation for 5 minutes in ice. After washing in FACS buffer (2% FBS, 2 mM EDTA in PBS, 0.2 µm-filtered), cell were resuspended in FcR block (1:200) and incubated on ice for 10 minutes. An equal amount of 2X antibodies was then added to each sample to reach the desired concentration. 2X mixed antibodies were prepared in brilliant dye buffer (BD Biosciences), whereas the single antibody was prepared in FACS buffer for reference controls. For blood monocytes, CD45 BV605 (BD Biosciences 563053), CD11b Pacific Blue (Biolegend 101223), and Ly6C BV570 (Biolegend 128029) antibodies were used at final dilutions of 1:500, 1:1000, and 1:1000, respectively.

Stromal cellular fraction (SVF) was collected from mice epididymal fat pads by enzymatic (1 mg/mL Collagenase D, Roche 1108888200) and mechanical (37°C, 160 rpm, 30 minutes) digestion, followed by removal of floating adipocytes from 100 µm-filtered and centrifuged cell suspension (600 xg, 5 minutes, 4°C). The remaining red blood cells in SVF were lysed in 500 µL RBC lysis buffer, incubation for 2-3 minutes in room temperature. Fc receptor was blocked and antibodies were added as described above for blood samples. For tissue macrophages, CD45 BV605 and F4/80 BV650 (Biolegend, 123149) were used, both at a final dilutions of 1:500.

After 20 min incubation with antibody on ice, free antibodies were washed off using FACS buffer and the cells were analyzed with the Cytek Northern lights 3L. The voltages of SSC and FSC channels were set to arbitrary unit of 100 and 25, respectively that were previously set for identification of blood lymphocytes. The stopping criteria was set to 50,000 single events. Finally, the signal for mixed antibodies were unmixed using the single reference controls from respective sample types, and background extracted using unstained samples. Data was further analyzed using FlowJo 10.10, and blood monocytes were gated for CD45^+^CD11b^+^Ly6C^+^ cells and tissue macrophages were gated for CD45^+^F4/80^+^ cells and data was represented as % of positive cells/single cells.

For detection of halo in immune cells, SVFs isolated from eWAT, sWAT, or BAT of mice on a chow or HFD diet were first stained with a viability dye (Zombie NIR; Biolegend, 423105, 1:1000 in PBS) on ice for 5 minutes. After wash in FACS buffer, the cells were resuspended in complete culture media supplemented with TMR direct halo ligand (G299A; Promega, 1:2000) for 20 minutes on ice. Cells were then washed with FACS buffer, pelleted, and re-suspended in 25 μL of Fc block (BD Biosciences, 553141, 1:100 in brilliant stain buffer) for a 15-minute incubation on ice. Cells were then stained with fluorophore-conjugated primary antibodies by directing adding 25 μL of a 2X antibody cocktail, which includes rat anti-mouse SiglecF-BV421 (clone E50-2440, BD Horizon, 562681), rat anti-mouse/human CD11b-Pacific Blue (clone M1/70, BioLegend, 101224), rat anti-mouse Ly6C-BV570 (clone HK1.4, BioLegend, 128030), rat anti-mouse F4/80-BV650 (clone BM8, BioLegend, 123149), rat anti-mouse Ly6G-BV711 (clone 1A8, BioLegend, 127643), rat anti-mouse/human B220-BV750 (clone RA3-6B2, BioLegend, 103261), Amenian hamster anti-mouse TCRβ-Alexa Fluor 488 (clone H57-597, BioLegend, 109215), rat anti-mouse CD45-PerCP (clone 30-F11, BioLegend, 103130), mouse anti-mouse CD64-PE/Dazzle594 (clone X54-5/7.1, BioLegend, 139320), rat anti-mouse PDGFRα-PE/Cy5 (clone APA5, BioLegend, 135920), Armenian hamster anti-mouse CD11c-PE/Cy5.5 (clone N418, Invitrogen/eBioscience, 35-0114-82), rat anti-mouse MHC-II-PE/Fire 810 (clone M5/114.15.2, BioLegend, 107667), rat anti-mouse CD19-Spark NIR 685 (clone 6D5, BioLegend, 115568), and mouse anti-mouse NK1.1-APC/Fire750 (clone PK136, BioLegend, 108752). After incubation on ice for 30 minutes, cells were washed with FACS buffer twice, and the final pellets were resuspended in 200 μL FACS buffer. 100 μL of the cell suspension were analyzed live on a Cytek Aurora spectral flow cytometer.

In addition, flow cytometry was done from whole blood cells of mice 5, 10 and 30 minutes after injecting with CFSE-labeled adipoEVs, and samples were processed as described above. In this study, the CFSE signal was measured in CD45^+^CD11b^+^Ly6C^+^ cells to assess adipoEV uptake by blood monocytes.

### Isolation of Mitochondria

Adipose tissue was homogenized in ice cold isolation buffer (10 mM MOPS, 1 mM EDTA, 210 mM mannitol, and 70 mM sucrose, pH 7.4) using a Potter-Elvehjem homogenizer. The homogenate was centrifuged for 5 minutes at 600 xg, 4°C. The resulting supernatant was filtered through cheese cloth and centrifuged for 15 minutes at 10,000 xg to pellet mitochondria. Mitochondria were resuspended in 0.02 µm-filtered PBS for flow cytometry analysis.

### Isolation of stromal vascular cells (SVF) and differentiation of adipocytes from SVF

Subcutaneous white adipose tissue (sWAT) from 4-6 week-old mice was digested for 1 hour at 37°C in buffer containing 100 mM HEPES, 120 mM NaCl, 50 mM KCl, 5 mM glucose, 1 mM calcium and 1 mg/ml collagenase D. The single cell suspension was filtered through a 100 µm cell strainer and centrifuged at 600 g for 5 minutes at 4°C. SVF cells were resuspended in culture media (DMEM/F12 media containing 10% FBS, GlutaMax, 1X penicillin-streptomycin and gentamicin) and filtered through a 45 µm cell strainer. Cells were centrifuged again as above. The cells were resuspended in culture media and grown at 37°C, 5% CO_2_. For in vitro differentiation, SVF cells (∼95% confluency) were treated with 500 µM 3-isobutyl-1-methylxanthine (IBMX), 1 µM dexamethasone, 5 µg/ml Insulin and 1 µm rosiglitazone for 2 days. Following the 2 days of adipogenic induction, cells were maintained in media containing only 5 µg/ml Insulin.

### Staining of EVs for in vivo injection

Adipocytes were cultured in FluoroBrite media (ThermoFisher) containing 1% exosome-depleted FBS (SBI: EXO-FBSHI-250A-1), 1X penicillin-streptomycin, 1X glutamax for 24 hours. Conditioned media was harvested and concentrated. Concentrated adipocyte conditioned media or plasma diluted 1:1 in PBS were combined with CFSE for a final concentration of 100µM. Samples were incubated at 37°C for 30 minutes and EVs purified by SEC. EVs were counted by nano particle tracking analysis (NTA; ZetaView by Particle Metrix) and 1×10^8^ labeled EVs per gram body weight (∼2×10^9^ EVs per 20g mouse) were injected.

### Adipose Tissue EV (ATEV) isolation

ATEVs were isolated as previously described^49^. Briefly, mice used for ATEV isolation were perfused with 3 ml ice-cold PBS through the left cardiac ventricle to remove blood from adipose tissues. Adipose tissue (∼1g) was harvested and digested as in the *Isolation of stromal vascular cells (SVF) and differentiation of adipocytes from SVF* section with the addition of 4mg/ml dispase. Once a single-cell suspension, the samples were centrifuged at 500xg for 5 minutes. The liquid between the floating adipocytes and pelleted SVF was harvested and centrifuged again at 500xg for 5 minutes to ensure all cells were removed. The sample was then centrifuged at 1200xg to remove apoptotic bodies. The total EV population was purified with size exclusion chromatography (10ml column; Cross-linked Sepharose CL-2B particles (Millipore Sigma). EVs were quantified by nano particle tracking analysis (NTA).

### Ex vivo incubation of adipose tissue

adipose tissues were harvested and ∼40mg was cultured in 300µl FluoroBrite media (ThermoFisher) containing 1% exosome-depleted FBS (SBI: EXO-FBSHI-250A-1), 1X penicillin-streptomycin, 1X glutamax for 6 hours. Media was harvested and centrifuged and EVs, purified as in the *Adipose Tissue EV (ATEV) isolation* section. EVs were quantified by nano particle tracking analysis (NTA).

### Immunogold staining and Electron Microscopy

Post perfusion fixation with 4% formaldehyde in phosphate buffered saline (PBS), samples were cut into 100 µm thick sections using a vibratome (Leica VT1200S, Vienna, Austria). Sections containing regions of interest were fixed again in 4% formaldehyde in PBS overnight at 4°C. Fixative was washed out of the sections with three 10 minutes rinses in PBS followed by incubation in fresh 0.5% sodium borohydride in PBS for 30 minutes to quench aldehydes. The samples were then rinsed 3 times in ultrapure water for 10 minutes each and incubated in 0.1% Triton X-100 in PBS for 30 minutes. After another 3x 10 minutes washes in ultrapure water, the samples were blocked in 10% normal goat serum (NGS), washed 3x 10 minutes in PBS, then incubated in primary antibody (anti-Halo; Promega G9281) diluted 1:500 with 1% bovine serum albumin (BSA) in PBS for 1 hour at room temperature, followed by 2 days at 4°C. Samples were washed 3X 10 minutes in PBS, and incubated in goat anti-rabbit 6nm diluted 1:200 with 1% BSA in PBS overnight at 4°C. Next, samples were washed 3x 10 minutes in PBS and incubated in either of the following;

A. 2% glutaraldehyde in PBS for 15 minutes, followed by 3x 10 minutes washes in PBS, then post fixed in 1% Osmium Tetroxide and 1.5% Potassium ferricyanide in PBS.
B. 2% glutaraldehyde, 0.05% ruthenium red, 0.2% tannic acid in PBS warmed to 37°C followed by 3x 10 minutes washes in PBS, then a mixture of 1% osmium, 1.5% potassium ferrocyanide, 0.05% ruthenium red in PBS on ice for 15-30 minutes.

After treating with A or B above, samples were washed with distilled water 3x 10 minutes then dehydrated with a graded acetone series 30%, 50%, 70%, 90%. 100% 3x for 10 minutes each step. Once dehydrated, samples were infiltrated with Spurr’s resin (Electron Microscopy Sciences, Hatfield, PA) 30%, 50%, 70%, 90%, 100% overnight, 2 100% spurs exchanges with microwave assistance, flat embedded and polymerized at 60°C for 48 hours. After curing, specific regions of interest were excised and mounted on a blank epoxy stub for sectioning. 70 nm sections were then cut, post-stained with 2% aqueous uranyl acetate and Sato’s lead and imaged on a TEM (JEOL JEM-1400 Plus) at an operating voltage of 120 kV.

### Statistical analysis

All data is presented as mean ± SEM. *p<0.05, ** p<0.01 and ***p<0.001 by two-tailed Student’s *t* test, or in the case of systemic assays two-way ANOVA. A p value < 0.05 was determined to be statistically significant. For all mouse studies, the *n* value corresponds to individual mice of a given treatment. Data was analyzed used Prism GraphPad Software.

## Acknowledgements

S.T., M.R., Y.L.P, and C.C. are supported by the National Institutes of Health (NIH; 1R21EB035738-0, 1R01DK137791), the DRC at Washington University (NIH P30DK020579), and the American Heart Association (AHA; 23IPA1054013). W.J. is supported by AHA 24POST1244220. This study was also supported by National Institutes of Health grants P30 DK056341 (Washington University Nutrition Obesity Research Center), UL1 TR000448 (Washington University School of Medicine Institute of Clinical and Translational Sciences), and DK137206. For electron microscopy, experiments and imaging were performed in part by the Washington University Center for Cellular Imaging (WUCCI) supported by Washington University School of Medicine, The Children’s Discovery Institute of Washington University and St. Louis Children’s Hospital (CDI-CORE-2015-505 and CDI-CORE-2019-813) and the Foundation for Barnes-Jewish Hospital (3770 and 4642). We thank the Alvin J. Siteman Cancer Center at Washington University School of Medicine and Barnes-Jewish Hospital in St. Louis, MO., for the use of the Siteman Flow Cytometry Core, which provided flow cytometry instrumentation. The Siteman Cancer Center is supported in part by an NCI Cancer Center Support Grant #P30 CA091842.

## Disclosures

SK serves on scientific advisory boards for Merck, Abbvie and Verdiva Bio.

