## Supplementary information for "Immune cells regulate circulating adipocyte extracellular vesicle levels in response to metabolic shifts"

PDF includes:

Figure S1-S4
Table S1-S3

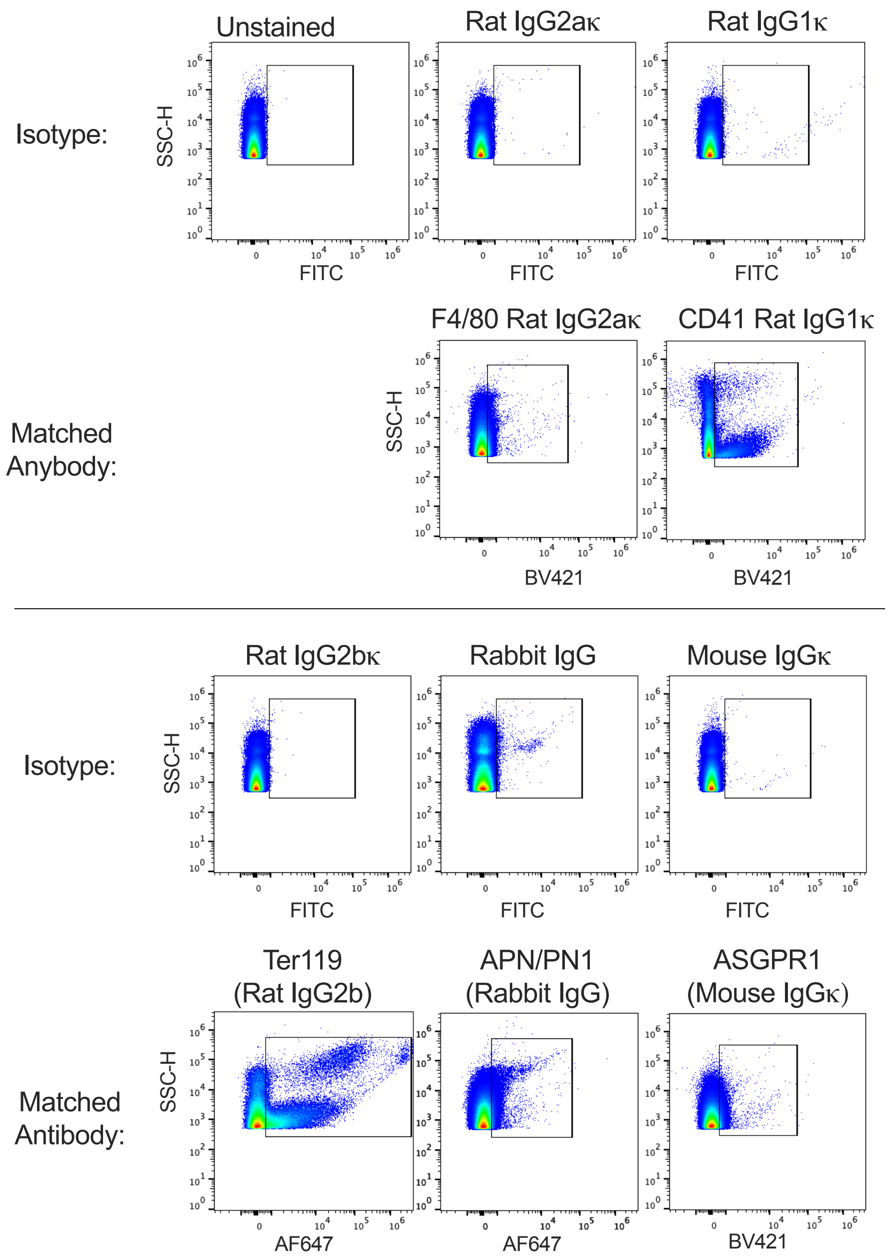

**Figure S1**. Isotype controls. Representative flow plots of antibodies used and their respective isotype control. All isotype control antibodies were used at the same concentration as the marker-specific antibodies.

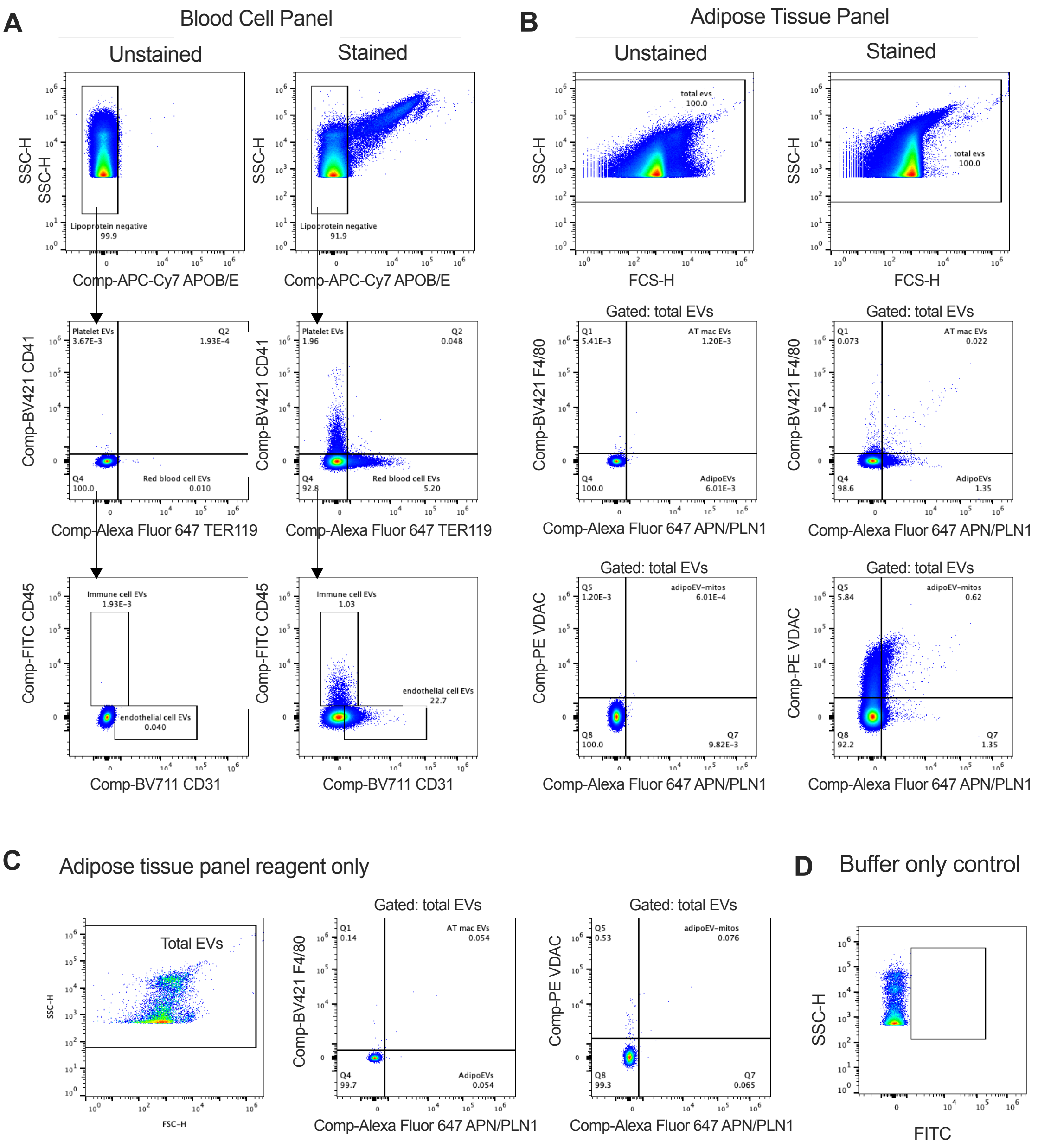

**Figure S2.** Gating strategy for EV panels. **A.** Plasma was stained for the blood cell panel (APPOB/E, TER119, CD31, CD41, and CD45). The lipoprotein-negative (APOB/E$^-$) population was gated for platelet (CD41$^+$/TER119$^-$) or red blood cell (TER119$^+$, CD41$^-$) EVs. The double negative population as gated for endothelial cell (CD31$^+$, CD45$^-$) or immune cell (CD45$^+$) EVs. The unstained control is shown. **B.** Plasma was stained for the adipose tissue panel (APN/PLN1, F4/80, and VDAC). Total plasma particles were gated for adipoEVs (APN/PLN$^+$, F4/80$^-$), or adipose tissue macrophage EVs (APN/PLN$^+$, F4/80$^+$). Total plasma particles were also plotted APN/PLN by VDAC to quantify total mitochondria (VDAC$^+$) and adipoEV-mitos (APN/PLN$^+$, VDAC$^+$). **C.** PBS combined with all reagents for the adipose tissue panel (antibodies, FcBlock, triton) at the concentrations used in B. Flow plots demonstrate the absence of fluorescence positive particles in this reagent only control. **D.** Events detected with buffer alone.

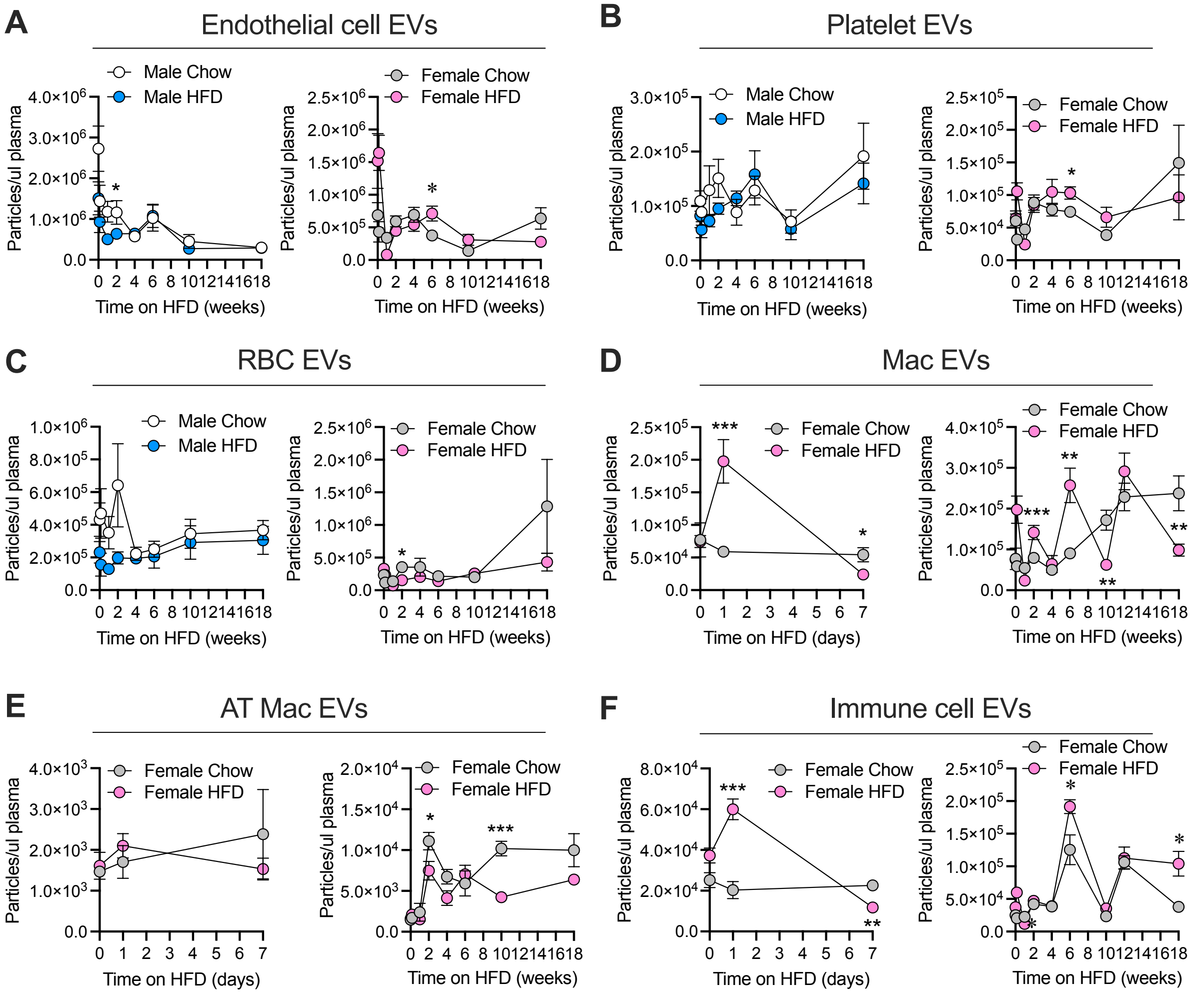

**Figure S3.** Blood cell and immune cell EV levels over a high-fat feeding time course. Wild type male or female mice were placed on a high-fat or matched low-fat diet and blood was collected at the indicated time on diet (N=6). Endothelial cell EVs ($CD31^+$, $CD45^-$; **A**), platelet EVs ($CD41^+$/$TER119^-$; **B**) or red blood cell EVs (RBC; $CD41^-$, $TER119^+$; **C**) were quantified. Females on a chow or high fat diet were bled at the indicated time points and total macrophage (Mac) EVs ($F4/80^+$; **D**) adipose tissue (AT) macrophage EVs ($APN/PLN^+$, $F4/80^+$; **E**), or total immune cell EVs ($CD45^+$; **F**), were quantified. Data are presented as mean ± s.e.m. *$P < 0.05$, ** $P < 0.01$, *** $P < 0.001$.

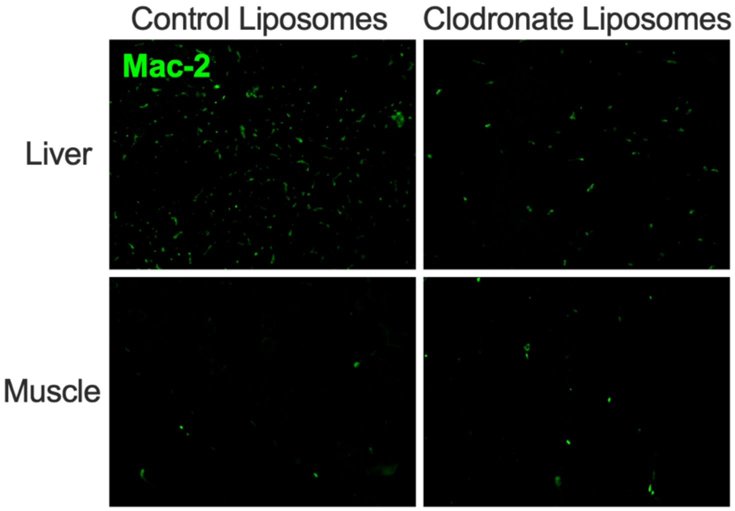

**Figure S4.** Macrophage content in liver and muscle with clodronate treatment. Liver or gastrocnemius muscle sections were immuno-stained with Mac-2 to label macrophages. Representative images of mac-2 stain in liver or muscle from control liposome-treated, or clodronate liposome treated mice.

**Table S1.** Estimated total circulating EVs from indicated cell types in mice. N=6-8.

| Cell type | Total EVs in a 20g mouse (~800ul plasma) | Fold change in obesity |
|---|---|---|
| Adipocyte | $5.7\text{x}10^{8} \pm 4.31\text{x}10^{8}$ | 3 |
| Myocyte | $3.7\text{x}10^{8} \pm 1.8\text{x}10^{8}$ | 1.2 |
| Hepatocyte | $5.0\text{x}10^{9} \pm 3.53\text{x}10^{9}$ | 2 |

**Table S2**. MIFlowCyt-EV check list

| Framework Criteria | What to report | Please complete each criterion | |
|---|---|---|---|
| 1.1 Preanalytical variables conforming to MISEV guidelines. | Preanalytical variables relating to EV sample including source, collection, isolation, storage, and any others relevant and available in the performed study. | Plasma EV samples were enriched by size exclusion chromatography | |
| 1.2 Experimental design according to MIFlowCyt guidelines. | EV-FC manuscripts should provide a brief description of the experimental aim, keywords, and variables for the performed FC experiment(s) using MIFlowCyt checklist criteria: 1.1, 1.2, and 1.3, respectively. Template found at www.evflowcytometry.org. | Reported in the manuscript | |
| 2.1 Sample staining details | State any steps relating to the staining of samples. Along with the method used for staining, provide relevant reagent descriptions as listed in MIFlowCyt guidelines (Section 2.4 Fluorescence Reagent(s) Descriptions). | Reported in the manuscript | |
| 2.2 Sample washing details | State any steps relating to the washing of samples. | EVs were purified away from free antibody, free dyes, and plasma proteins by size exclusion chromatography. | |
| 2.3 Sample dilution details | All methods and steps relating to sample dilution. | Reported in the manuscript | Samples were diluted with 0.02 nm-filtered PBS. |
| 3.1 Buffer alone controls. | State whether a buffer-only control was analyzed at the same settings and during the same experiment as the samples of interest. If utilized it is | 0.02 nm-filtered PBS. See Supplemental Figure S2 C. | |

| | | |
|---|---|---|
| | recommended that all samples be recorded for a consistent set period of time e.g. 5 minutes, rather than stopping analysis at a set recorded event count e.g. 100,000 events. This allows comparisons of total particle counts between controls and samples. | |
| 3.2 Buffer with reagent controls. | State whether a buffer with reagent control was analyzed at the same settings, same concentrations, and during the same experiment as the samples of interest. If used state what the results were. | Reagent only control included (Figure S2B Antibody mix was combined with 0.02nm-filtered PBS and processed by size exclusion chromatography like samples. |
| 3.3 Unstained controls. | State whether unstained control samples were analyzed at the same settings and during the same experiment as stained samples. If used, state what the results were, preferably in standard units. | Each experiment had an unstained control sample analyzed at the same time and same settings. This control was used for gating positive events. |
| 3.4 Isotype controls. | The use of isotype controls is applicable to immunofluorescence labelling only. State whether isotype controls were analyzed at the same settings and during the same experiment as stained samples. If utilized, state which antibody they are matched to, the concentration used, and what the results were (Section 4.2, 4.3, 4.4). Due to conjugation differences between manufacturers if should be stated if the isotype controls are from the same manufacturer as the matched antibodies. | Isotype controls are in Figure S1. All isotype controls were matched to the highest concentration of that specific isotype used. Isotype control samples were run with the same settings as samples stained with specific antibodies. See table S1 for isotypes and concentrations. |

| | | |
|---|---|---|
| 3.5 Single-stained controls. | State whether single-stained controls were included. If used state whether the single-stained controls were recorded using the same settings, dilutions, and during the same experiment as stained samples and state what the results were, preferably in standard units (Section 4.2, 4.3, 4.4). | Single stained controls were used for reference controls. Fluorescent antibodies were conjugated to FSP CompBeads (Cytek). Labeled beads were analyzed at the same fluorescence gain as EV samples. Antibody-bead ratio was optimized to ensure fluorescence was not saturated. |
| 3.6 Procedural controls. | State whether procedural controls were included. If used, state the procedure and if the procedural controls were acquired at the same settings and during the same experiment as stained samples. | The same mouse plasma sample was run each day with experimental samples to assess day-to-day variability. |
| 3.7 Serial dilutions. | State whether serial dilutions were performed on samples and note the dilution range and manner of testing. The fluorescence and/or scatter signal intensity would ideally be reported in standard units (see Section 4.3, 4.4) but arbitrary units can also be used. This data is best reported by plotting the recorded number events/concentration over a set period of time at different sample dilution. The median fluorescence intensity at each of the dilutions should also ideally be plotted on the same or a separate plot. | Samples were serially diluted to find the best concentration where EVs are detected above buffer background and that the abort rate was less than the 10 percent of the count rate to avoid swarming artifacts. $2.5 \times 10^9$ total particles (as quantified by nanoparticle tracking analysis) in the 50µl staining mix before dilution was found to give the best event rate to abort rate ratio. Due to variability of EV concentration between cohorts, the dilution for each experiment was altered to prevent swarming (1:2 to 1:5). The particle numbers were corrected for with the dilution factor. |
| 3.8. Detergent treated EV-samples | State whether samples were detergent treated to assess lability. If utilized, state what detergent was used, the end | EVs were not treated with detergent. |

| | | |
|---|---|---|
| | concentration of the detergent, and what the results were of the lysis. | |
| 4.1 Trigger Channel(s) and Threshold(s). | The trigger channel(s) and threshold(s) used for event detection. Preferably, the fluorescence calibration (Section 4.3) and/or scatter calibration (Section 4.4) should be used in order to report the trigger channel(s) and threshold(s) in standardized units. | 405 SSC trigger and the threshold set at 600 arbitrary units. |
| 4.2 Flow Rate / Volumetric quantification. | State if the flow rate was quantified/validated and if so, report the result and how they were obtained. | Samples were run at the lowest flow rate setting (~13µl/min). The internal flow rate sensor of the cytometer measured flow rate. |
| 4.3 Fluorescence Calibration. | State whether fluorescence calibration was implemented, and if so, report the materials and methods used, catalogue numbers, lot numbers, and supplied reference units for the standards. Fluorescence parameters may be reported in standardized units of MESF, ERF, or ABC beads. The type of regression used, and the resulting scatter plot of arbitrary data vs standard data for the reference particles should be supplied. | No fluorescence channels were calibrated. |
| 4.4 Light Scatter Calibration. | State whether and how light scatter calibration was implemented. Light scatter parameters may be reported in standardized units of nm2, along with information required to reproduce the model. | SSC detector was calibrated with Rosetta beads (Exometry). |
| 5.1 EV diameter/surface | State whether and how EV diameter, surface area, and/or volume has been | Rosetta beads (Exometry) beads were used to |

| | | |
|---|---|---|
| area/volume approximation. | calculated using FC measurements. | estimate EV size by Mei theory. |
| 5.2 EV refractive index approximation. | State whether the EV refractive index has been approximated and how this was done. | Not applicable |
| 5.3 EV epitope number approximation. | State whether EV epitope number has been approximated, and if so, how it was approximated. | Not applicable |
| 6.1 Completion of MIFlowCyt checklist. | Complete MIFlowCyt checklist criteria 1 to 4 using the MIFlowCyt guidelines. Template found at www.evflowcytometry.org. | Complete |
| 6.2 Calibrated channel detection range | If fluorescence or scatter calibration has been carried out, authors should state whether the upper and lower limits of a calibrated detection channel were calculated in standardized units. This can be done by converting the arbitrary unit scale to a calibrated scaled, as discussed in Section 4.3 and 4.4, and providing the highest unit on this scale and the lowest detectable unit above the unstained population. The lowest unit at which a population is deemed ‘positive’ can be determined a variety of ways, including reporting the 99th percentile measurement unit of the unstained population for fluorescence. The chosen method for determining at what unit an event was deemed positive should be clearly outlined. | Done |
| 6.3 EV number/concentration. | State whether EV number/concentration has been reported. If calculated, it is preferable to report EV | Concentration were reported |

| | | |
|---|---|---|
| | number/concentration in a standardized manner, stating the number/concentration between a set detection range. | |
| 6.4 EV brightness. | When applicable, state the method by which the brightness of EVs is reported in standardized units of scatter and/or fluorescence. | Not applicable |
| 7.1. Sharing of data to a public repository. | Provide a link to the experimental data in a public data repository. | |

**Table S3.** Antibodies for EV flow cytometry

| Antibody | Catalogue # | Dilution | Final antibody concentration in sample (pg/µl) |
|---|---|---|---|
| CD31 anti-mouse BV711 Clone MEC13.3 | BD Biosciences 740680 | 1:500 | 0.4 |
| CD31 anti-human Clone WM59 BV 711 | BioLegend 303135 | 1:8000 | 0.006 |
| CD45 Rat Anti-Mouse FITC Clone 30-F11 | BD Biosciences 553080 | 1:200 | 2.5 |
| CD45 anti-human BV421 Clone HI30 | BioLegend 304031 | 1:250 | 0.1 |
| CD41 anti-mouse BV421 Clone MWReg30 | BioLegend 133911 | 1:200 | 1 |
| TER-119 anti-mouse AF647 Clone TER-119 | BioLegend 116218 | 1:200 | 2.5 |
| F4/80 anti-mouse BV421 Clone BM8 | BioLegend 123131 | 1:200 | 1 |
| Anti-VDAC1/Porin + VDAC2 antibody Anti- mouse/human PE | Abcam ab317896 | 1:5000 | 0.1 |
| Adiponectin anti-mouse/human CL647 | ThermoFisher PA1-054 | 1:100 | 2.85 |
| Perilipin-1 anti-mouse/human AF647 | Cell Signaling 52975 | 1:100 | Concentration not reported |
| CD29 (ITGB1) anti-mouse PE/Cyanine7 Clone HMβ1-1 | BioLegend 102221 | 1:200 | 1 |
| CD63 anti-mouse PE/Cyanine7 Clone NVG-2 | BioLegend 143909 | 1:1000 | 0.2 |

| | | | |
|---|---|---|---|
| CD9 anti-mouse PE/Cyanine7 Clone MZ3 | BioLegend 124815 | 1:1000 | 0.2 |
| ASGPR-1 mouse BV421 anti-mouse/human Clone 8D7 | BD Biosciences 742697 | 1:1000 | 0.2 |
| Myosin Heavy Chain (MHC) ant-mouse/human PE Clone MF20 | BD Biosciences 564408 | 1:200 | 1 |
